# Natural selection maintains species despite widespread hybridization in the desert shrub *Encelia*

**DOI:** 10.1101/2020.01.23.917021

**Authors:** Christopher D DiVittorio, Sonal Singhal, Adam B Roddy, Felipe Zapata, David D Ackerly, Bruce G Baldwin, Craig R Brodersen, Alberto Burquez, Paul VA Fine, Mayra Padilla Flores, Elizabeth Solis, Jaime Morales-Villavicencio, David Morales-Arce, Don W Kyhos

## Abstract

Natural selection is an important driver of genetic and phenotypic differentiation between species. A powerful way to test the role of natural selection in the formation and maintenance of species is to study species complexes in which potential gene flow is high but realized gene flow is low. For a recent radiation of New World desert shrubs (*Encelia*: Asteraceae), we use fine-scale geographic sampling and population genomics to determine patterns of gene flow across two hybrid zones formed between two independent pairs of species with parapatric distributions. After finding evidence for extremely strong selection at both hybrid zones, we use a combination of field experiments, high-resolution imaging, and physiological measurements to determine the ecological basis for selection at one of the hybrid zones. Our results identify multiple ecological mechanisms of selection (drought, salinity, herbivory, and burial) that together are sufficient to maintain species boundaries despite high rates of hybridization. Given that multiple pairs of species hybridize at ecologically divergent parapatric boundaries in the adaptive radiation of *Encelia*, such mechanisms may maintain species boundaries throughout this group.

**SIGNIFICANCE STATEMENT:** In Baja California, the deserts meet the coastal dunes in a narrow transition visible even from satellite images. We study two species pairs of desert shrubs (Encelia) that occur across this transition. Although these species can interbreed, they remain distinct. Using a combination of genetics, field experiments, 3D-imaging, and physiological measurements, we show that natural selection counteracts the homogenizing effects of gene exchange. The different habitats of these species create multiple mechanisms of selection - drought, salinity, herbivory, and burial, which together maintain these species in their native habitats and their hybrids in intermediate habitats. This study illustrates how environmental factors influence traits and fitness and how this in turn maintain species, highlighting the importance of natural selection in speciation.

## INTRODUCTION

A major goal in biology is to understand the balance between natural selection and gene flow in speciation. While natural selection is believed to play a role in the formation of most species, under what conditions it alone can generate and maintain phenotypic and genetic differences in the presence of high rates of gene flow is unknown (1–3). According to the theory of ecological speciation, adaptation to different habitats and tradeoffs in resource use and allocation can generate strong divergent natural selection, manifested as low fitness of migrant and hybrid phenotypes (1, 4). However, despite the appeal of adaptation as a robust mechanism of species formation and maintenance, many population genetic models of divergence with gene flow require levels of natural selection that may be unrealistically high or uncommon in nature (5–7). To accommodate this discrepancy, many speciation models require prezygotic barriers to evolve quickly after adaptation has been initiated in order to “complete” speciation by locking in locally adaptive traits (7, 8). For example, prezygotic isolating mechanisms such as assortative mating, shifts in phenology, and geographic isolation are frequently invoked as operating alongside postzygotic isolating mechanisms, such as selection against hybrids, presumably because prezygotic sources of isolation are thought to be difficult to reverse (9–11). Some systems, however, do not fit this paradigm and species remain distinct despite exhibiting little evidence for assortative mating and little geographic or phenological isolation. Such systems are ideal for testing the potential for divergent natural selection to maintain species boundaries because they exhibit high rates of hybridization (potential gene flow), but low rates of introgression (realized gene flow) (12, 13).

The radiation of desert shrubs in the genus *Encelia* (Asteraceae) is a powerful system for investigating ecological mechanisms of species formation and maintenance. The eighteen described *Encelia* species and subspecies are morphologically and physiologically diverse and occupy a variety of specialized edaphic and climatic niches (14–16). Still, no evidence of intrinsic barriers to reproduction has been found in experimental crosses between different taxon pairs in the genus (16, 17). Additionally, hybrid zones are found between eight unique species pairs (18), yet these zones are almost always narrowly limited to ecotones or zones of disturbance, a pattern known to be indicative of extrinsic or ecological control over hybrid zone structure (1, 19). Moreover, the same pollinators are frequently observed moving between sympatric and parapatric species (16, 17). As with most desert plants, flowering occurs primarily in response to rainfall, synchronizing flowering phenology at regional scales (20). Thus, there are few potential sources of prezygotic isolation to explain the maintenance of species boundaries at the numerous known hybrid zones.

The hybrid zones between *Encelia palmeri* - *E. ventorum* (*palmeri* - *ventorum*) and *E. asperifolia* - *E. ventorum* (*asperifolia* - *ventorum*) illustrate these patterns particularly well. These species are endemic to the Baja California Peninsula of México where *E. ventorum* inhabits coastal sand dunes and *E. palmeri* and *E. asperifolia* occupy inland desert plains (Figure 1). Across both sets of hybridizing taxa, species exhibit morphological divergence that putatively reflects adaptation to their contrasting environments. Although species are strictly limited to their respective habitats, hybrid zones form at the linear ecotones between dune and desert habitats (Figure S1). For example, previous data from one *palmeri* - *ventorum* hybrid zone showed that hybrids are narrowly limited to the ecotone where their abundance frequently approaches that of the parental species and that the interspecific hybridization rate for plants ∼200 m from the ecotone is 3.6% (16). These hybrids are vigorous and produce large quantities of viable propagules, yet parental taxa have maintained adjacent distributions in some locations for at least 125 years without fusing (16, 21). Based on these observations, we hypothesized that strong natural selection maintains adaptive divergence despite high rates of hybridization. We tested this hypothesis by asking the following three questions: (a) What is the magnitude and direction of hybridization and introgression between taxa in each of the two hybrid zones? Then, for the *palmeri* - *ventorum* hybrid zone, we further ask: (b) What is the magnitude and direction of phenotypic selection in dune, desert, and ecotone habitats? (c) What are the ecological mechanisms generating selection on hybrid and parental phenotypes?

**Figure 1:**
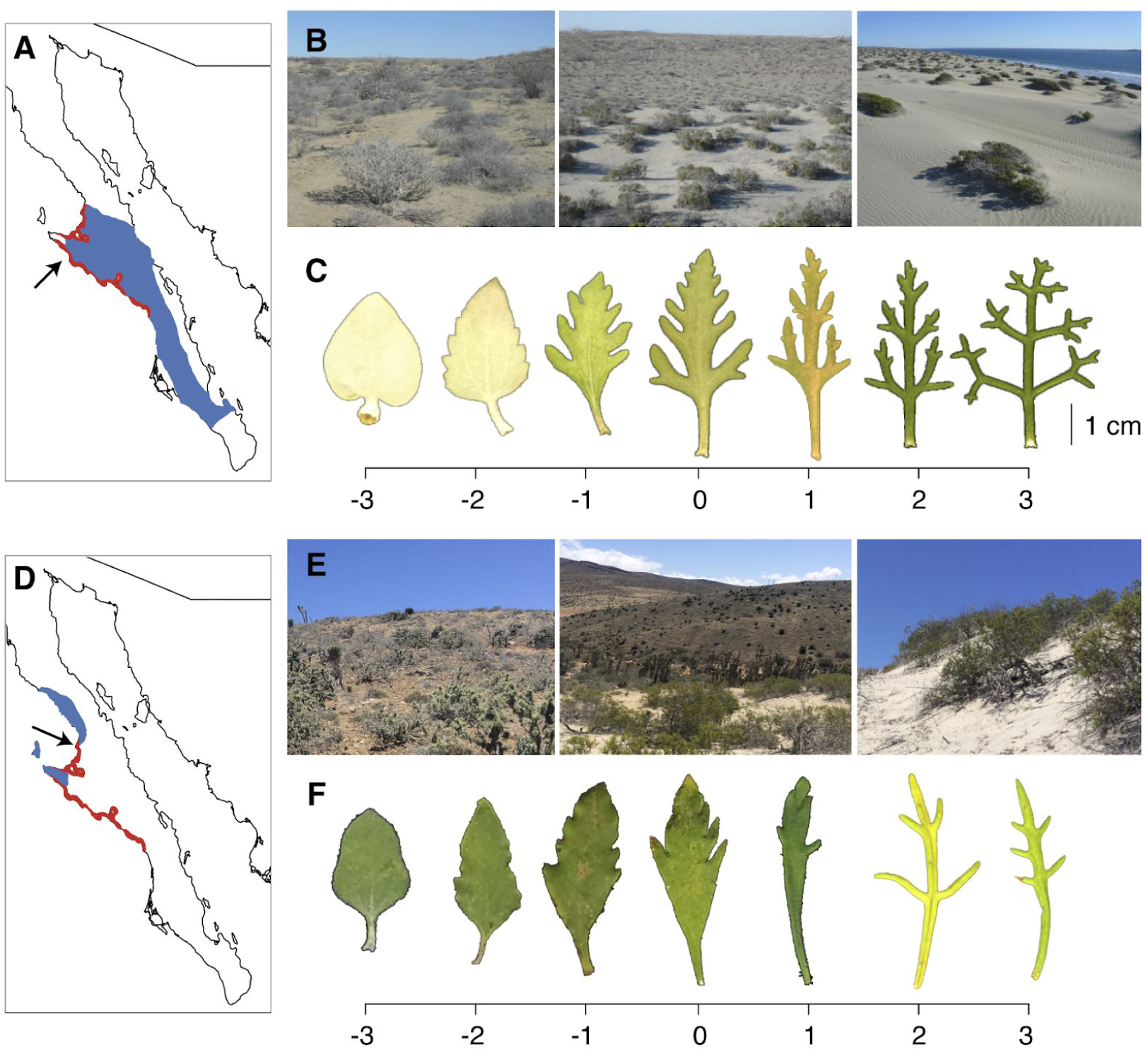
(a) Map of Baja California Sur, showing the range of *Encelia palmeri* (blue) and *E. ventorum* (red) with the San Roque experimental site marked by the arrow. (b) Photographs of (L - R) desert, ecotone, and dune habitats at San Roque. (c) Representative leaf phenotypes of *E. palmeri* (L), *E. ventorum* (R), and hybrids (center). Scale bar at bottom is a multivariate hybrid index composed of the first axis of a principal components analysis of leaf shape and area. (d) The range of *E. asperifolia* (blue) and *E. ventorum* (red) with the Punta Lobos sampling site marked by the arrow. (e) Photographs of (L - R) desert, ecotone, and dune habitats at Punta Lobos. (f) Representative leaf phenotypes of *E. asperifolia* (L), *E. ventorum* (R), and hybrids (center).

## RESULTS

### Patterns of gene flow

#### Hybridization, or potential gene flow

To estimate the potential for gene flow between *palmeri* - *ventorum* and *asperifolia* - *ventorum*, we genotyped and phenotyped 112 and 91 adult individuals, respectively, across parental habitats and the ecotone transition. Across both taxon comparisons, genetic and morphological indices of admixture were highly correlated (*r* = 0.95, pval <2e-16 and *r* = 0.94, pval <2e-16 for *palmeri* - *ventorum* and *asperifolia* - *ventorum*, respectively; Figure 2). Whether hybrids were diagnosed using genetic or morphological measures, we found that 72% (N = 54 for *palmeri* - *ventorum*) and 61% (N = 38 for *asperifolia* - *ventorum*) of plants in the ecotones were hybrids (Figure S2). In addition, for *palmeri* - *ventorum*, we estimated the rate of present-day hybridization. To do so, we sampled fruits from naturally occurring, phenotypically-pure parental plants located within ∼200 m of either side of the ecotone, germinated these fruit seeds in a greenhouse, and measured the percentage of offspring that were hybrids. We found that 31.4% (*n* = 175) of the progeny of *E. palmeri* and 5.6% (*n* = 409) of the progeny of *E. ventorum* showed evidence of interspecific hybridization or backcrossing (see *Methods: Estimates of Current Hybridization*). The abundance of hybrids in the hybrid zone and the large proportion of hybrid seed outside of the hybrid zone suggest potential gene flow into and across the hybrid zone is high.

**Figure 2:**
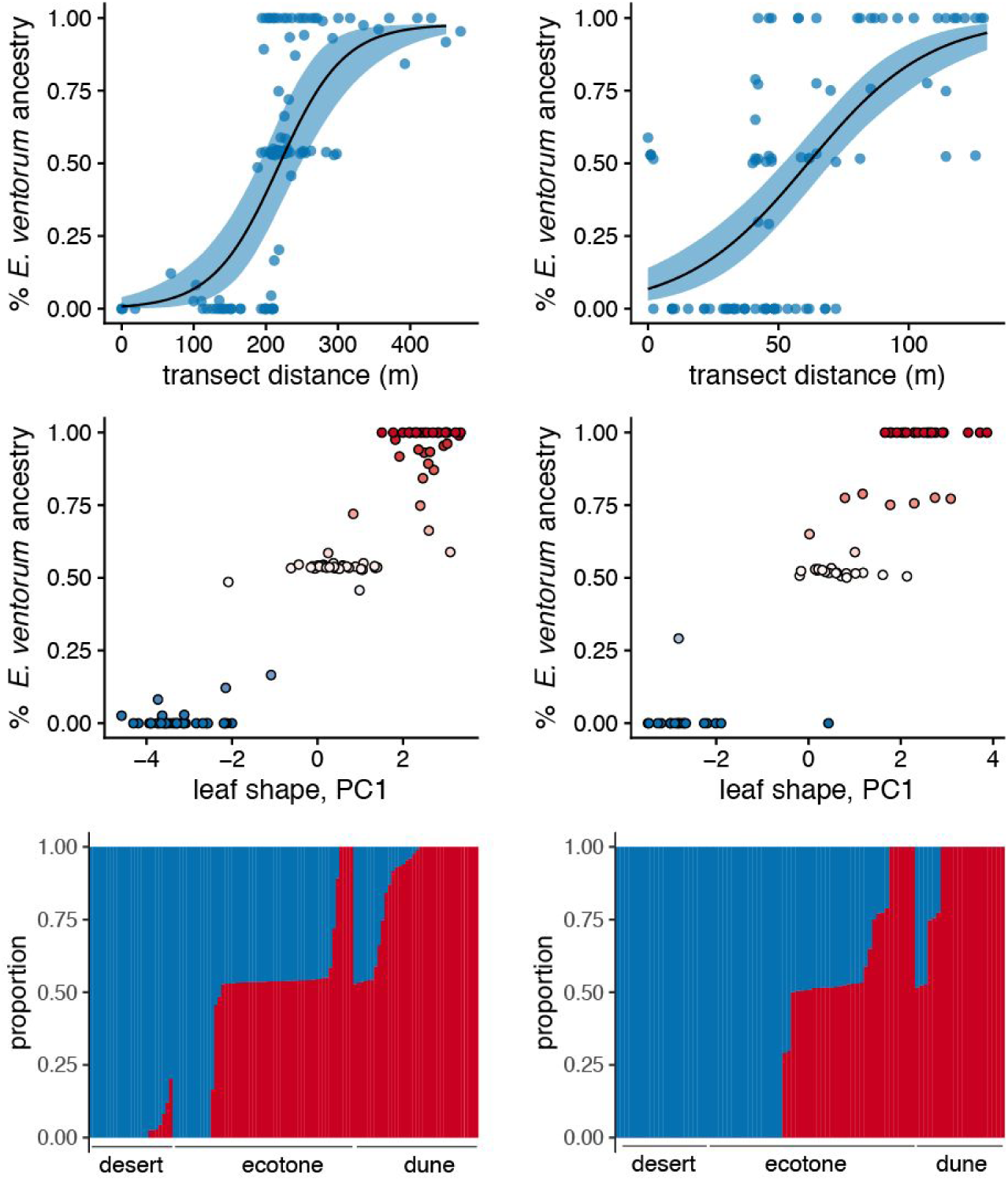
Patterns of hybridization and introgression at two natural hybrid zones in *Encelia*. Shown left is *E. palmeri* - *E. ventorum* (N = 112 individuals); right is *E. asperifolia* - *E. ventorum* (N = 91). Shown top to bottom: inference of cline width based on genomic ancestry estimates, the distribution of individuals in phenotypic and genetic space, and inference of genomic ancestry for individuals in the hybrid zone based on variant data from ddRAD loci. Ancestry results are shown ordered by habitat and then ancestry proportion; blue is *E. palmeri* (L) or *E. asperifolia* (R) and red is *E. ventorum*. Across both hybrid zones, we see narrow clines given effective dispersal of these species, limited introgression beyond the hybrid zone, and evidence that most hybrids are F1s. Together, these results suggest extremely strong selection is structuring these hybrid zones and thus also maintaining species boundaries between these hybridizing pairs.

Our results do not exclude the possibility that some forms of prezygotic isolation exist between these species. Despite this, our results show the potential for gene flow exists beyond the hybrid zone, at rates that would lead to species fusion if not counteracted by natural selection.

#### Introgression, or realized gene flow

In contrast to potential gene flow, we inferred low levels of realized gene flow between *palmeri* - *ventorum* and *asperifolia* - *ventorum*. Using the hybrid zone phenotypic and genetic data, we characterized the extent of introgression through both hybrid zones. Across both hybrid zones, we found similar patterns: narrow clines coincident with habitat transitions, limited evidence for introgression beyond the ecotone, and an abundance of F1 hybrids (Figure 2). Cline widths were estimated as 118 m and 93 m for *palmeri* - *ventorum* and *asperifolia* - *ventorum*, respectively. This discrepancy between potential and realized gene flow suggests divergent natural selection prevents introgression outside of the ecotone, thus maintaining species boundaries despite rampant hybridization. Indeed, for the *palmeri* - *ventorum* hybrid zone, the hybrid zone coincides with a steep gradient in soil type and with a near-complete turnover of the composition of the rest of the perennial plant assemblage (Figure 3). Given the overlap between the hybrid zone and the habitat transition, we hypothesized that habitat-mediated selection is likely structuring this hybrid zone.

**Figure 3:**
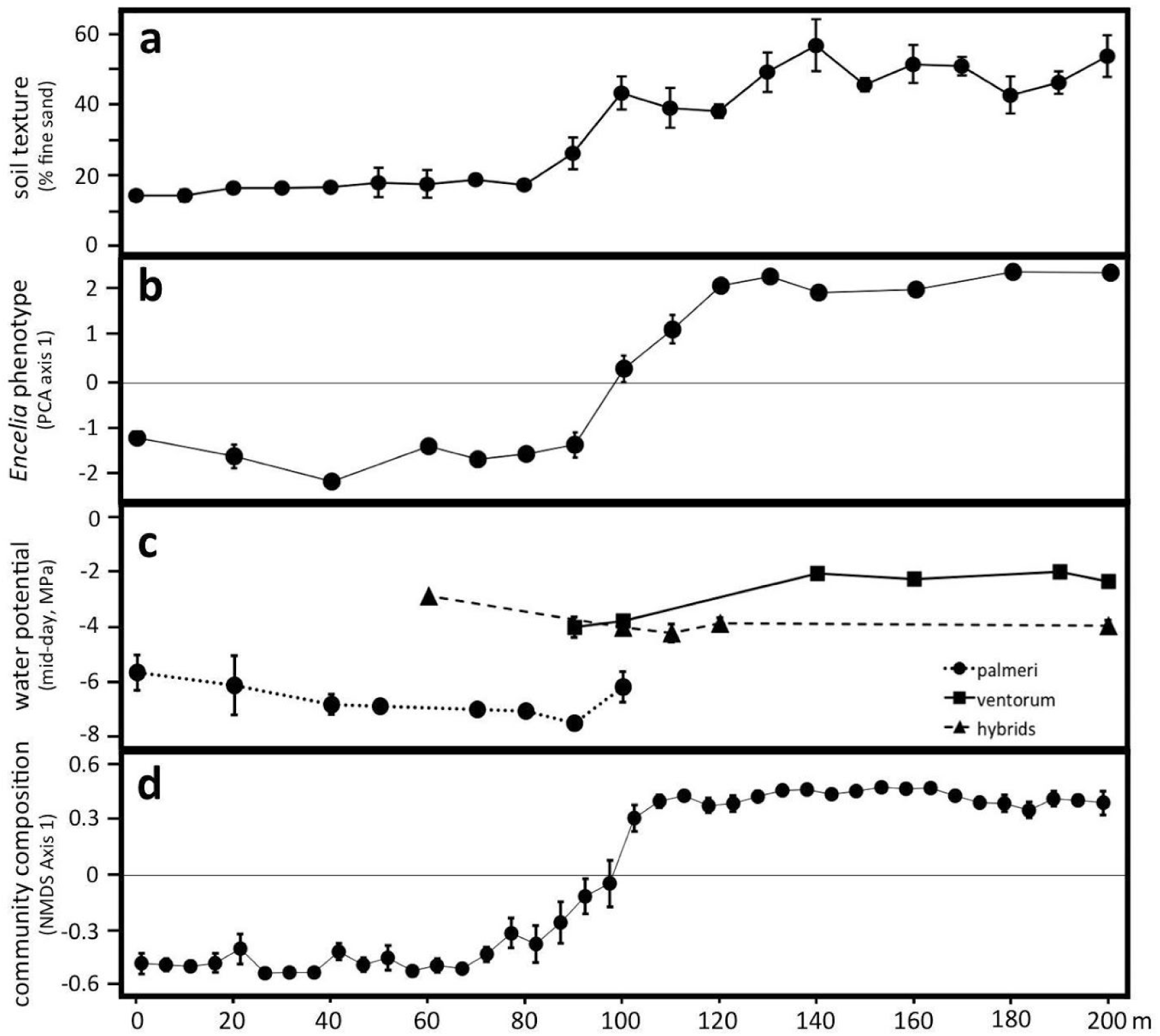
Transects centered on and perpendicular to the ecotone at the San Roque hybrid zone reveal concordant clines in (a) soil texture, (b) *Encelia* leaf morphology, (c) *Encelia* mid-day stem water potential, and (d) species composition of the rest of the perennial plant assemblage. The transect begins in the desert (*E. palmeri* habitat) and ends in the dune (*E. ventorum* habitat). Measurements were performed along seven replicate 200 meter transects. Error bars are ± one standard error.

### Patterns of natural selection

To test the hypothesis that strong habitat-mediated selection is preventing introgression beyond the ecotone, we conducted a reciprocal transplant field experiment in one of the two hybrid zones: the *palmeri* - *ventorum* hybrid zone at the San Roque experimental site. This field-based experiment allowed us to measure the direction and magnitude of natural selection acting in dune, ecotone, and desert habitats (*Methods: Reciprocal Transplant*; Figure S1). Parental taxa and hybrids were grown from seed, collected and transplanted into dune, ecotone, and desert habitats, watered for two months, and then allowed to grow naturally for five more months. Aboveground and belowground biomass, survival, and reproduction were assayed at the end of the experiment. This experiment revealed extremely strong divergent natural selection (Figure 4), which is on par with some of the strongest natural selection measured between naturally hybridizing taxa (22–25). Significant phenotype by habitat interactions were found for growth, survival, and composite fitness (Table S1), with selection coefficients against parental migrants ranging from *s* = 0.755 for *E. ventorum* in the desert to *s* = 0.983 for *E. palmeri* in the dunes. Hybrids were selected against in both parental habitats, with selection coefficients of *s* = 0.702 in the dune habitat and *s* = 0.097 in the desert habitat. In the ecotone habitat, hybrids outperformed the parental species (*E. palmeri*; *s* = 0.582 and *E. ventorum*; *s* = 0.460), although this difference was not statistically significant. Taken together, the results from the transplant experiment are consistent with the hypothesis that natural selection prevents introgression beyond the ecotone.

**Figure 4:**
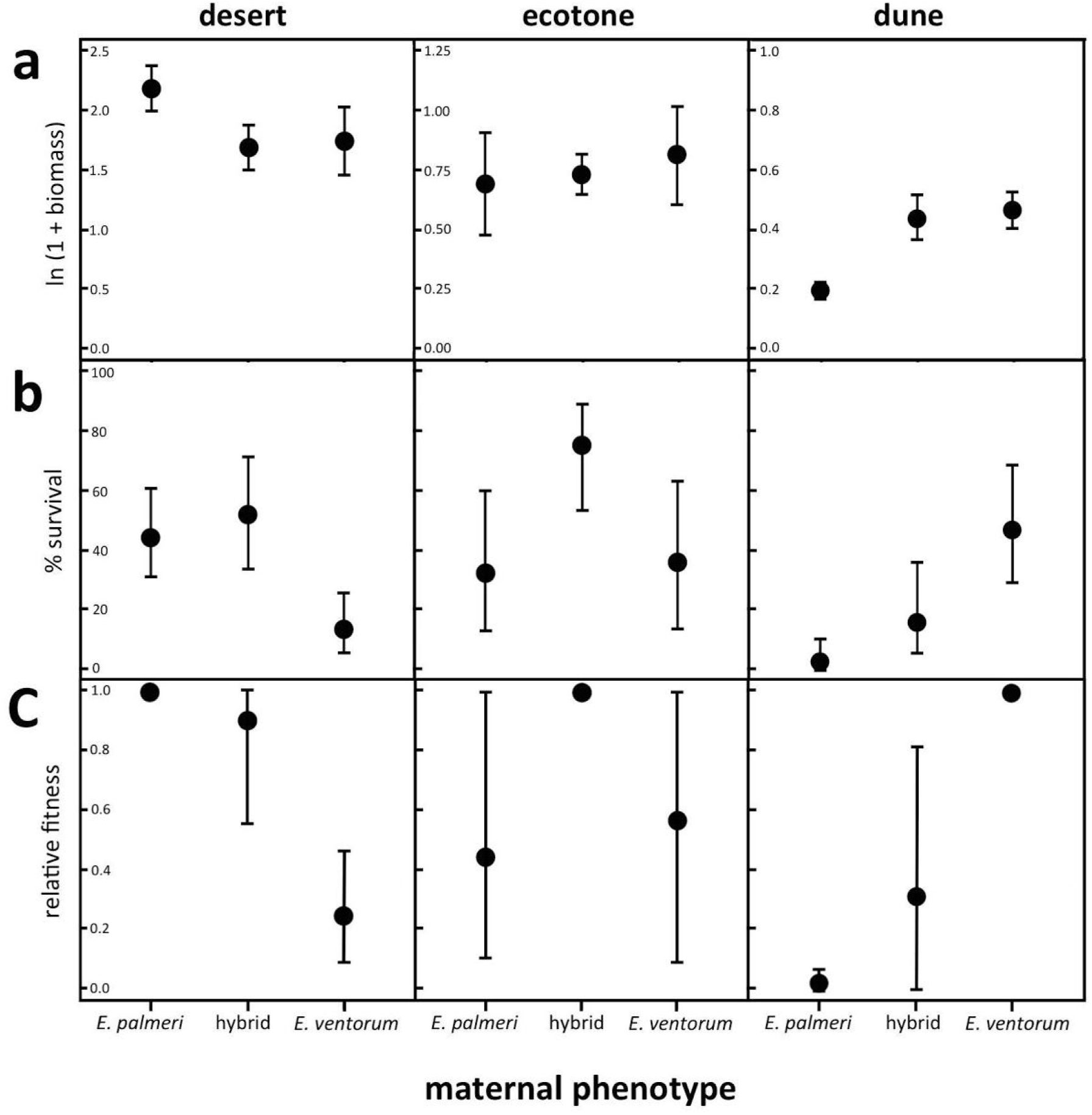
Results of the reciprocal transplant field experiment illustrating (a) growth measured as total ln-transformed biomass, (b) percent survival to the end of the experiment, and (c) composite relative fitness obtained by multiplying growth and proportional survival and setting the most fit phenotype in each habitat equal to one. Error bars for biomass are ± one standard error, error bars for survival are binomial 90% confidence intervals, and error bars for relative fitness are bootstrapped 95th percentile confidence intervals truncated to between 0 and 1 (*Methods: Reciprocal Transplant*).

### Mechanisms of natural selection

The desert and dune environments differ in water availability, wind strength, soil salinity, and herbivory pressure, which we hypothesized creates selection gradients between the two habitats and contributes to the patterns of selection measured in the reciprocal transplant experiment.

#### Water availability

The desert habitat is drier, hotter, and less humid than the dune habitat (Figures S3-S4). The coarser texture of dune sand results in more rapid water infiltration, allowing precipitation from fog, dew, and small rain events to percolate deeper than in adjacent desert soils (Figure S3) (26). Once water infiltrates it is strongly insulated from evaporation, creating a persistent lens of water suspended within the dune that is a common feature of both coastal and inland dune systems (27, 28).

We experimentally tested the importance of this gradient in water availability by adding water to a subset of the plants in the reciprocal transplant experiment (*Methods: Resource Addition*). Increasing water availability equalized fitness differences in the desert habitat, eliminating divergent natural selection (Figure 5). By contrast, adding water had no effect on growth or survival of either species in the dune habitat (Table S1). These results show that the gradient in water availability between habitats is necessary but not sufficient to cause divergent natural selection.

**Figure 5:**
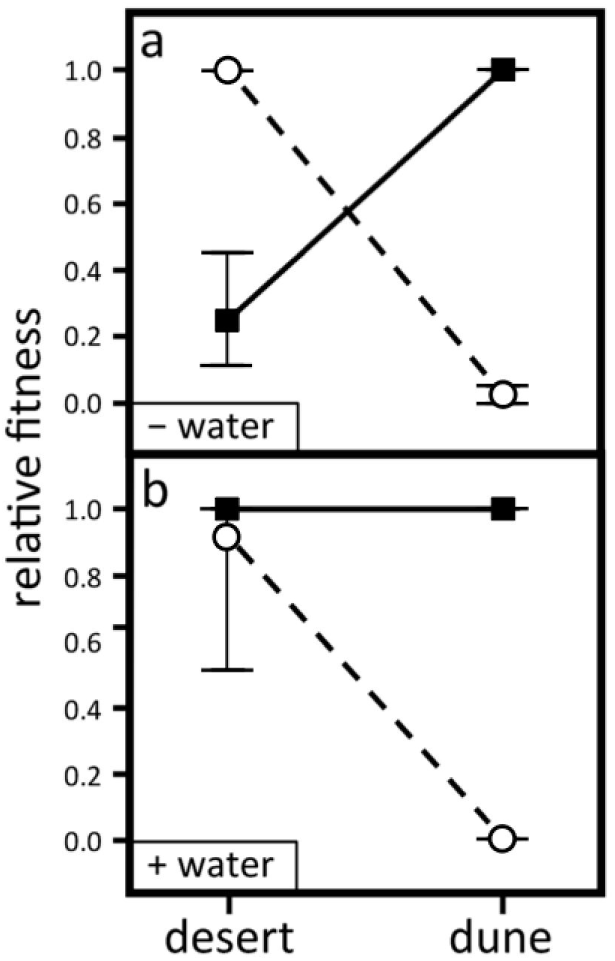
Relative fitness of *E. palmeri* (circles) and *E. ventorum* (squares) in dune and desert habitats, (a) without supplemental water, and (b) with supplemental water. Relative fitness is calculated proportionally, with the most fit taxon in each habitat set to one. Error bars are bootstrapped 95th percentile confidence intervals truncated to between 0 and 1.

Hydraulic traits of *E. palmeri, E. ventorum*, and hybrids differed in ways that are consistent with published studies on plant adaptation to drought (Supplemental *Methods: Hydraulic Physiology*). Xylem vessel diameter, stem hydraulic conductance, leaf hydraulic capacitance, and mid-day stem water potential were all predicted to be lower in *E. palmeri* from xeric desert habitats than *E. ventorum* from mesic dune habitats (29, 30). Xylem vessel diameter varied in the predicted directions with vessel diameter smallest in *E. palmeri*, largest in *E. ventorum*, and intermediate in hybrids (Figure S5; Table S2; one-way ANOVA: *p* = 0.013, *F* = 4.36, *df* = 2). Leaf hydraulic capacitance also varied in the predicted directions, with capacitance lowest in *E. palmeri*, greatest in *E. ventorum*, and intermediate in hybrids (Figure S6). Mid-day stem water potential at the center of the hybrid zone also varied in the predicted directions, with *E. palmeri* exhibiting more negative water potentials than hybrids or *E. ventorum* (Figure 3c; one-way ANOVA: *p* < 0.001, *F* = 23.4, *df* = 27). Stem hydraulic conductance was the only trait that did not vary in the predicted direction, with *E. palmeri* exhibiting greater conductance per unit leaf area than *E. ventorum* (Table S2; two-sample t-test: *p* = 0.046, *F* = 2.61, *df* = 5). However, because conductance is needed to supply water to leaves, conductance may be higher where atmospheric demand is greater (31), such as in the drier air of *E. palmeri* habitat (Figure S4).

#### Wind

The dunes inhabited by *E. ventorum* are created by strong onshore winds that transport beach sand inland (32, 33); the specific epithet *ventorum* translates as “blowing winds”. These winds create stable ecotones between dune and desert habitats defined by the tangent of the direction of the wind to the curvature of the beach (Figure S1) (34, 35). A variety of biomechanical stresses are generated by wind including abrasion of the leaf cuticle and structural failure from flapping (36). Additionally, wind results in burial and excavation of whole plants as dunes shift vertically and laterally (37). We tested the importance of wind as a component of natural selection by scoring mortality due to burial in the reciprocal transplant experiment (*Methods: Reciprocal Transplant*). A plant was scored as killed by burial if it was completely covered with sediment, exhibited dry or necrotic tissues when excavated, and had been healthy at previous time points (Figure S7). Analysis of mortality due to burial revealed a significant habitat-by-phenotype interaction (Table S1). Mortality from burial was highest in the dunes (22.2%), less in the ecotone (7.0%), and not observed in the desert habitat (0%). Across all three habitats, mortality from burial only affected *E. palmeri* (17.5%) and hybrids (9.0%), with no individuals of *E. ventorum* dying from burial during the course of the experiment (Figure S8).

Several morphological traits unique to *E. ventorum* may function as adaptations to wind and burial. First, *E. ventorum* exhibits heavily dissected, glabrous leaves (Figure 1), traits that in other species reduce aerodynamic drag and dampen the flapping instability that can cause structural failure (36). Second, *E. ventorum* exhibits pronounced juvenile apical dominance relative to *E. palmeri* resulting in a narrow, conical canopy architecture (Figure S9) that is known to function as an adaptation to avoid burial (37, 38). Finally, unlike other species in the genus, *E. ventorum* exhibits adventitious rooting, which is known to function as an adaptation to track shifting sand dunes (37).

#### Salinity

Salt spray is a major stressor in many coastal habitats (39). To test for a gradient in salt stress we first quantified variation in the degree of leaf succulence, a known response to salinity that functions to dilute the internal salt concentration of leaves (40). Using leaf thickness as a proxy for succulence, we measured the thickness of leaves along a transect between dune and desert habitats, and also around the circumference of a large *E. ventorum* individual growing on the exposed foredune (*Supplemental Methods: Hydraulic Physiology*). Consistent with our predictions, leaf thickness was two to three times greater in the foredune habitat than near the desert (Figure S10b), and was also two to three times greater on the windward, ocean-exposed side of the canopy compared to the leeward side (Figure S10c).

To further investigate whether these patterns in leaf succulence corresponded to a gradient of salt stress, we measured leaf osmotic potential of the storage parenchyma cells in *E. ventorum* leaves along a transect from desert to dune. We hypothesized that leaf osmotic potential would become more negative with increasing proximity to the ocean because salt is passively taken up by the roots and because salt deposition on the leaves is a function of exposure to onshore winds (40, 41). Leaf osmotic potential was measured by excising and macerating internal non-photosynthetic parenchymous tissue under a saturated atmosphere and measuring the water potential of the macerated tissue using thermocouple psychrometers (*Supplemental Methods: Hydraulic Physiology*). This method eliminates remaining xylem tension and turgor pressure, so that the residual water potential is equal to the osmotic potential. Consistent with our hypothesis, leaf osmotic potential decreased from the desert to the foredune, indicating that salt stress increases with proximity towards the ocean (Figure S10a; linear regression: *p* < 0.01, *R*^2^ = 0.80, *n* = 8).

Additional evidence for the prevalence of salt stress comes from three-dimensional X-ray computed microtomographic (microCT) reconstructions of the stems of parental taxa and hybrids, which revealed the presence of irregular crystals inside and around resin ducts (Figure 6b). These crystals were abundant in *E. ventorum*, rare in *E. palmeri*, and present in intermediate amounts in hybrids (Table S2; one-way ANOVA: *p* < 0.001, *F* = 18.09, df = 2). Repeated microCT imaging of the stems as they desiccated revealed that these ducts are normally hydrated and that the crystals form as the stem desiccates. Since active salt sequestration in stems is a known adaptation to salinity in other taxa (40, 41), we hypothesized that these crystals were condensed ocean-derived salts.

**Figure 6:**
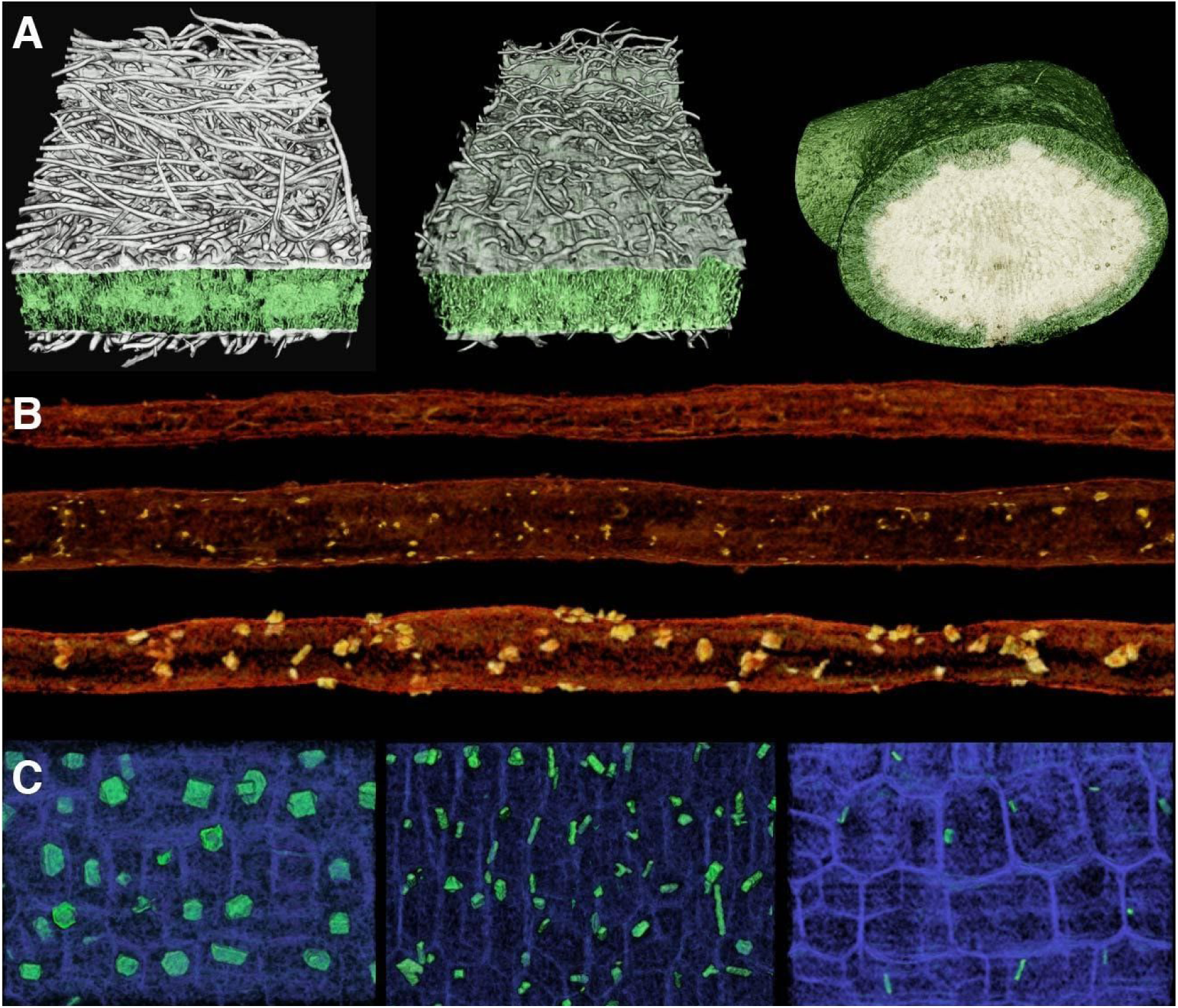
(a) Three-dimensional oblique-transverse images of leaf laminas of (left to right) *E. palmeri*, a hybrid, and *E. ventorum*, obtained using microCT imaging. In *E. ventorum*, the lightly colored cells in the center of the leaf are storage parenchyma holding salt. Exposed transverse surfaces are 820 µm, 820 µm, and 2000 µm wide, respectively. (b) Representative three-dimensional microCT images of resin ducts of (top to bottom) *E. palmeri*, a hybrid, and *E. ventorum*, showing differences in size and abundance of associated crystals. Resin ducts shown are all 1 mm long. (c) Representative microCT images of stem pith cells (blue) of (left to right) *E. palmeri*, a hybrid, and *E. ventorum*, highlighting differences in the size, shape, and abundance of pith crystals (green). All images are 300 µm by 200 µm.

To test this hypothesis, we measured the X-ray mass attenuation coefficients (MACs) of crystals from parental taxa and hybrids, and compared them to the MACs of a number of candidate compounds (Table S3). The crystals associated with resin ducts had average MACs ranging from 7.16 to 7.51 placing them within the range of the MACs of the solid forms of the major salt constituents of seawater (NaCl = 6.26, MgCl_2_ = 7.95, KCl = 10.12). In comparison, several chromenes and benzofurans, known constituents of the resins in *Encelia* (42), have MACs less than 0.2. Other compounds were also poor matches, including silicon dioxide (SiO_2_ = 2.73) and calcium oxalate (CaC_2_O_4_ = 5.17), two secondary compounds frequently found in crystalline form in plants (43).

#### Herbivory

A variety of herbivores were observed consuming *Encelia* plants at the San Roque experimental site. To determine whether herbivory contributed to divergent selective pressures we first analyzed sources of mortality in the reciprocal transplant experiment (Figure S8). Plants were diagnosed as having been killed by herbivory if they were completely defoliated, if tracks or other signs of herbivore activity were found nearby, and if the plant was otherwise healthy at previous time points (*Methods: Reciprocal Transplant*). Analysis of patterns of herbivory and mortality indicated a significant habitat-by-phenotype interaction (Table S1). In the desert habitat, *E. ventorum* was the most heavily consumed (23%), followed by hybrids (8%), and *E. palmeri* (2%). In the dune habitat *E. palmeri* and *E. ventorum* both sustained 14% mortality from herbivory while no hybrids were killed by herbivory.

Additionally, due to the higher resource availability in the dune habitat and the higher growth rate of *E. ventorum*, we predicted that *E. ventorum* would exhibit lower investment in secondary compounds and structures than *E. palmeri* due to a tradeoff between investment in growth versus herbivore defense (44). One conspicuous difference between *E. ventorum* and *E. palmeri* is the presence of a dense pubescence of trichomes on the leaves of *E. palmeri* and hybrids, which are known to function as a mechanical defense against herbivory (45). Although trichomes may have other functions, including reflecting solar radiation and increasing boundary layer thickness (29), the density of trichomes varied in the directions predicted based on patterns of herbivory (Figure 6a; *E. palmeri* = 447.6 ± 9.1 mm^-2^, hybrids = 84.6 ± 13.0, *E. ventorum* = 0.17 ± 0.17; one-way ANOVA: *p* < 0.001, *F* = 224.7, df = 2).

*Encelia* species are also known for their diversity of secondary compounds, many of which are known defenses against herbivory (42, 46). In *E. palmeri*, microCT analysis revealed that the pith cells of the stems contained a large number of conspicuous crystals that differed in shape and location from those previously identified in the resin ducts (Figure 6c). These crystals were most abundant in *E. palmeri*, less frequent and smaller in hybrids, and very infrequent and smallest in *E. ventorum* (Table S2; one-way ANOVA on abundance: *p* < 0.001, *F* = 217.1, df = 2). The MACs of these crystals (*E. palmeri* = 5.17 ± 0.15, hybrids = 5.32 ± 0.11, *E. ventorum* = 5.19 ± 0.11) closely matched the MAC of calcium oxalate (CaC_2_O_4_ = 5.17), a secondary compound known to act as a mechanical and chemical deterrent to herbivory in a large number of plant taxa (43). Together these results indicate that herbivory likely contributes to divergent selective pressures, but that further study is needed to disentangle the multiple sources of selection acting on secondary compounds and structures.

## DISCUSSION

Using a combination of fine-scale genetic and phenotypic characterization of hybrid zones, manipulative field experiments, three-dimensional anatomical characterizations, and ecophysiological measurements, we show how ecologically-based divergent natural selection maintains phenotypic and genetic divergence between co-distributed species despite widespread hybridization. Our work further demonstrates how strong ecological tradeoffs can be created through multiple agents of selection. In particular, we demonstrate that collinear gradients of water, burial, salinity and herbivory all contribute to divergent natural selection. Moreover, while the net effect of all sources of selection was divergent, no single factor alone exhibited divergent selective pressures. Rather, each factor either operated strongly in one habitat but was neutral in the other, or operated in the same direction in both habitats but stronger in one. Although we have yet to identify the ecological mechanisms of selection acting in the *asperifolia* - *ventorum* hybrid zone, the similarities between the two hybrid zones suggests similar processes are structuring this hybrid zone as well (Figure 2). Thus, *Encelia* builds on other studies that show that ecologically divergent evolutionary lineages maintained by selection may be more common than is appreciated, especially if adaptation involves adjustments in traits that are hard to measure or not readily visible, and involves adaptation to multiple ecological gradients simultaneously (4, 47–50).

These results further show that selection acts throughout the life cycle of these organisms, against both parental migrants and hybrids. If parental seeds disperse into the wrong habitat, they are unlikely to survive (Fig. 4), leading to parental species largely being restricted to their native habitat and reducing the frequency of hybridization (Fig. S2; (4, 6, 13, 51)). Pollen still disperses easily across habitat boundaries, however, leading to 5 - 30% of seeds in each parental habitat being admixed. Despite this high potential for gene flow, hybrids are still selected against relative to native parental phenotypes (Fig. 4), strongly limiting realized gene flow (Fig. 2). Thus, as found in other studies (6, 52), these results show the same ecological mechanisms were responsible for selection against both parental migrants and hybrids.

Given that the species that we studied are not fully reproductively isolated and that selection appears to be maintaining species boundaries, what is the likely fate of these species? These species might be at the early stages of divergence for which additional prezygotic isolating barriers have not yet evolved by reinforcement (53). However, the evolution of reinforcement is less likely in systems in which the cost of hybridization is low. Because individual plants produce thousands of seeds with low levels of parental investment, selection for the evolution of reinforcement is likely to be weak ((54); but see (55)).

Nevertheless, given enough divergence time, hybrid incompatibilities should evolve, even as taxa hybridize (56). Yet, until additional barriers to reproduction evolve, the mechanisms maintaining species boundaries in this system appear entirely driven by environmental selection. As a corollary, if this environmental gradient erodes due to changing conditions, the boundaries between these species might erode as well (57). In fact, field-based observations have found hybrid swarms and later-generation backcrosses mainly in disturbed habitats like arroyos and roadsides (16).

In sum, our findings show how natural selection can be an effective barrier to introgression, even when hybridization rates are high. Our study further provides explicit ecological mechanisms for four different selective agents agents (water availability, salinity, burial, and herbivory), underscoring that adaptation to different habitats likely involves complex interactions between multiple agents of selection acting on whole phenotypes. Natural hybrid zones associated with habitat transitions can also be found between eight species pairs in *Encelia* (18), yet no cases of phenotypic fusion or species collapse are known. Our study thus provides a mechanistic basis for the rapid generation of diversity in *Encelia* as it radiated throughout the deserts of North and South America, and showcases the potential power of natural selection in generating and maintaining biodiversity.

## METHODS

### Overview

We first used fine-scale phenotypic and genotypic sampling to determine how hybridization patterns change across narrow ecotones between two pairs of desert plant species, *Encelia palmeri* - *E. ventorum* and *E. asperifolia* - *E. ventorum*. Concordant patterns across both hybrid zones justified focus on one pair: *palmeri* - *ventorum*. For this pair, we used a reciprocal transplant experiment to test the hypothesis that strong natural selection was structuring the hybrid zone. After supporting our hypothesis, we conducted fine-scale analyses of the anatomy and physiology of *palmeri* - *ventorum* to determine possible mechanisms of selection. Full details on our methods are available in the Supplementary Information.

### Site Characteristics

All sampling, field measurements, and experiments for *palmeri* - *ventorum* were carried out at San Roque, a settlement on the Pacific Coast of Baja California Sur, México (Figure 1). All sampling for *asperifolia* - *ventorum* were carried out on Punta Lobos, a site on the Pacific Coast of Baja California Sur, México (Figure 1). Both sites are characteristic of the sand dune-desert ecotones found throughout the Baja California Peninsula. Afternoon winds across the peninsula are consistent and predictable (32, 33, 58) (Figure S11), which creates stable, linear ecotones between dune and desert habitats with dunes moving parallel to the ecotone (Figure S1). Habitat type can be identified by eye based on differences in soil color and texture that arise due to differential sand content between the desert and the dune (Figure 1b, 1e); desert soil comprises ∼15% fine sand whereas dune soil comprises ∼55% fine sand (Figure 3). The ecotone is defined as the narrow 20 m linear region at the interface between these habitats (Figure 3a).

### Hybrid zone analysis

To analyze patterns of hybridization and introgression at the *palmeri* - *ventorum* and *asperifolia* - *ventorum* hybrid zones, we analyzed phenotypic and genetic data from adult plants across transects in each hybrid zone. For each individual, we sampled multiple adult leaves and recorded both latitude and longitude and habitat type. We followed a linear transect through the hybrid zone, starting in the desert and ending in the dune. In the ecotone, we sampled perpendicular to the transect due to the narrowness of the transition. In total, we sampled 112 individuals for the *palmeri* - *ventorum* hybrid zone and 91 individuals for the *asperifolia* - *ventorum* hybrid zone.

To characterize genetic patterns at the hybrid zones, we collected double-digest restriction aided (ddRAD) data from each individual. Per hybrid zone, we assembled loci and called variants across all individuals using Velvet (59), vsearch v2.4.3 (60), bwa v0.7.17 (61) and ANGSD v0.923 (62). To determine patterns of admixture and introgression across individuals, we used NGSadmix v32 to calculate ancestry proportions for each individual (63). We used HZAR to estimate genetic clines based on ancestry proportions (64).

To collect and analyze phenotypic data from the hybrid zones, we measured leaf area and shape by photographing leaves with a scale bar using a digital camera and then analyzing the images in Image-J (U.S. National Institutes of Health, http://imagej.nih.gov/ij/). After conducting a scaled and centered PCA on all leaves from a given hybrid zone, we averaged PC scores per individual across all leaves measured for that individual. PC1 captured 52% of the variation in the *asperifolia* - *ventorum* hybrid zone and 45% of the variation in the *palmeri* - *ventorum* hybrid zone. In both hybrid zones, PC1 largely reflected leaf shape (Figure 1C and 1F).

### Estimates of current hybridization

To obtain current, field-based estimates of hybridization rates in *E. palmeri* and *E. ventorum*, we collected fruits (cypselae) from phenotypically pure individuals growing near the hybrid zone and assayed their progeny for traits indicative of interspecific hybridization. In 2016, we collected cypselae from several hundred *E. palmeri* and *E. ventorum* mothers and germinated them in indoor flats filled with potting soil. After germination, any seedling that visually scored between −1 and 1 on the hybrid index (Figure 1C) was recorded as hybrid.

### Environmental characterization of the *palmeri* - *ventorum* hybrid zone

We characterized the abiotic and biotic conditions across the dune-desert ecotone in *E. palmeri* - *E. ventorum*. To measure microclimates, we used paired weather stations and soil moisture arrays to measure relative humidity, temperature, wind speed, leaf wetness, and soil moisture (Figure S3, 4). In the dune, we additionally measured photosynthetic photon flux density and wind direction. To measure soil texture, we characterized the percent of soil that was “fine sand” across 20 equidistant points across a 200m long transect (Figure 3a). To characterize biotic conditions, we performed community surveys of all non-*Encelia* vascular perennial plant species across seven 10 × 200 m belt transects. We summarized the resulting community data matrix using the “vegdist” function in the “vegan” package, and the “isoMDS” function in the “mass” package in R (Figure 3d).

### Reciprocal Transplant

We conducted a reciprocal transplant field experiment during 2010 - 2011 at the San Roque experimental site. In fall 2010, 276 parental and hybrid plants grown from seed were planted into dune, desert, and ecotone habitats. We measured growth, survival, and reproduction over the course of one growing season. In March 2011, all surviving plants were harvested and measured for plant size (stem basal diameter, plant height, canopy diameter, aboveground and belowground biomass), leaf morphology (leaf color, thickness, area, and shape), and (when possible) mid-day shoot water potential (see *Hydraulic Physiology*).

Growth was measured as total above-plus belowground biomass at the end of the experiment, or the total biomass of the plant at the time of its death. Survival was calculated as the percent of plants that were alive at the end of the experiment. Only one plant from the water addition experiment flowered during the course of the experiment, so reproductive output was not analyzed. Growth and survival were additionally combined into a composite fitness measure by multiplying a binary (0, 1) survival score with the ln(1+*x*) transformed biomass. This composite measure is a reasonable approximation of fitness, given that total plant size is an important predictor of seed production (14). Relative fitnesses were calculated by standardizing by the phenotype with the highest fitness (Figure 4c). Negative selection coefficients were calculated as one minus the relative fitness (24).

To test for the effects of habitat and phenotype on fitness, we constructed a general linear model using a negative binomial variance and link function implemented with the “glm.nb” function in R on ln(1+*x*) transformed data.

### Water Addition

We tested the hypothesis that water availability affected relative fitness of *E. palmeri* and *E. ventorum* by adding supplemental water to a subset of the plants in the reciprocal transplant field experiment. Five plants of *E. palmeri* and eight plants of *E. ventorum* were randomly chosen in each of dune and desert habitats to continue receiving water for an additional three months after an initial two-month watering period. Following completion of the experiment, plants were harvested and assayed for biomass and leaf traits. To test the effects of the watering treatment on growth, we constructed a general linear model using a negative binomial link function using the “glm.nb” function in R, first specifying a model with a water × phenotype interaction, and then specifying a model with water and phenotype independently added. Model fit was compared through a likelihood ratio (LR) test.

### Sources of mortality

We surveyed all experimental reciprocal transplant plants two or three times a week to record instances of mortality due to herbivory and from burial by wind-blown sand. Herbivory was scored if the entire plant was destroyed between visits, with evidence of herbivore activity nearby such as tracks or the herbivore itself, and no other proximate cause identified (e.g. bad weather). Mortality due to burial was scored if the entire plant was completely buried by sand and if all leaves were wilted and lacking turgor when excavated (Figure S7).

To analyze patterns of burial in the reciprocal transplant experiment, we performed backwards stepwise log-linear analysis on three-way contingency tables. To test for a habitat-by-phenotype interaction, we constructed models as described for the ‘Water addition’ experiment and performed LR tests on the two best fitting models (Table S1). We repeated this analysis for patterns of herbivory.

### Hydraulic Physiology

To characterize physiological differences between *E. palmeri* and *E. ventorum*, we measured five aspects of plant hydraulic physiology: leaf hydraulic capacitance, stem hydraulic conductance, mid-day shoot water potential, leaf succulence, and leaf osmotic potential. We focused on hydraulic physiology because the reciprocal transplant experiment suggested that water availability imposes a significant selection gradient on these taxa. Mid-day shoot water potentials were measured on mature, naturally occurring plants, whose locations were mapped using GPS and binned according to their distance from the hybrid zone. Water potentials were measured using a Scholander-type pressure chamber (PMS, Albany, Oregon; Model 1000). All measurements were made between 13:00 and 15:00 hours on sunny days. One healthy ten-centimeter long shoot per plant was excised using a razor blade and immediately placed into the pressure chamber. Water potentials of distance-binned individuals were averaged with error bars representing one standard error (Figure 3C). Differences in water potential at the ecotone habitat were analyzed using one-way analysis of variance (ANOVA). Detailed methods for other measurements of hydraulic physiology are available in the Supplemental Methods.

### X-ray Microtomography

To characterize divergence in fine-scale leaf and stem anatomy, we obtained high-resolution, three-dimensional (3D) images of stem and leaf structure by performing hard X-ray computed microtomography (microCT) at the Advanced Light Source, Lawrence Berkeley National Laboratory (LBNL), Beamline 8.3.2 (65, 66). Stem samples were obtained by collecting cuttings of *E. palmeri*, *E ventorum*, and hybrids growing naturally in the field in March 2015 and allowing them to slowly desiccate. Leaf samples were collected from plants growing in cultivation at the Agricultural Operations Station, University of California, Riverside in June 2015 and transported to LBNL, keeping the leaves hydrated until analysis. Samples were placed on a rotating stage in the 24 keV synchrotron X-ray beam and 1025 two-dimensional projections were recorded as the sample rotated continuously from 0 to 180 degrees. These raw tomographic projections were assembled using Octopus 8.7 (University of Ghent, Belgium) and then analyzed in Avizo 8.1 software. The resolution of the resulting images was 1.24 µm and 1.36 µm for stems and leaves, respectively.

For analysis of crystal structure in stems, we used 16-bit reconstructions. In Octopus 8.7, the mass attenuation coefficient (MAC) was calculated by manually drawing a line transect through fifteen crystals of each morphological type in each taxon and averaging peak pixel values. Mean and standard error for each crystal type in each phenotype were calculated for comparison to predicted MAC values for candidate crystal compounds obtained from an online database maintained by Argonne National Laboratory (http://11bm.xray.aps.anl.gov/absorb/absorb.php). We then surveyed the literature to build a list of candidate compounds and queried this database for predicted MAC values of each candidate compound. This list of candidate compounds included the most abundant salts in seawater, several secondary compounds previously found in *Encelia* resins, including benzofurans and benzopyrans, and other compounds commonly occurring in crystalized forms in plants (Table S3).

## Data accessibility

● Code used in genomic analysis: https://github.com/singhal/encelia
● Raw ddRAD reads: SRA (TBD)
● Individual genotypes and phenotypes from each hybrid zone: DataDryad (TBD)

## Acknowledgements

Funding for CTD was provided by the University of California Institute for México and the United States (UC-MEXUS), Consejo Nacional de Ciencia y Tecnología de México, US NSF (DEB-1011606), Heckard Endowment of the Jepson Herbarium, ARCS Foundation, California Desert Research Fund, Jiji Foundation, California Botanical Society, California Native Plant Society, Terra Peninsular A.C., American Genetics Association. SS was supported by a US NSF Postdoctoral Research Fellowship in Biology (#1519732). Field and laboratory assistance were provided by Maria Elide Arce-Aguilar, Lynette Butsuda, Benjamin Carter, Justin Chin, Kenneth DiVittorio, Melissa Ferriter, Emma Liffick, Camilla Morales-Arce, Meno Morales-Arce, Edna Morales-Arce, Ricardo Pereira, and Sarah Richman. Logistical support was provided by Juanita and Lenin Morales, John and Dianne Trotter, Juan and Shari Arce, Ejido Heroes de Chapultepec, and the Delegación de Bahía Asunción. Helpful discussions about previous versions were provided by Exequiel Ezcurra, Ivone Giffard-Mena, Rosemary Gillespie, Jorge Montiel, Ricardo Pereira, Daniel Rabosky, Montgomery Slatkin, Wayne Sousa, Sula Vanderplank, and David Wake. MicroCT imaging was performed at the Lawrence Berkeley National Laboratory Advanced Light Source, Beamline 8.3.2, supported by the Director, Office of Science, Office of Basic Energy Sciences, U.S. Department of Energy (DE-AC02-05CH11231).

## Author Contributions

CD, SS, AR, DA, PF, BB, CB, AB, DK designed research. CD, SS, AR, JM, DM, MPF, ES performed work. CD, SS, AR, FZ analyzed data and wrote the paper.

## Materials and Correspondence

All requests for materials should be made to the corresponding authors, Christopher DiVittorio and Sonal Singhal.

## References

1. D. Schluter, The Ecology of Adaptive Radiation (Oxford University Press, 2000).

2. L. H. Rieseberg, A. Widmer, A. M. Arntz, J. M. Burke, Directional selection is the primary cause of phenotypic diversification. Proc. Natl. Acad. Sci. U. S. A. 99, 12242–12245 (2002).

3. J. A. Coyne, H. A. Orr, Speciation (Sinauer Associates, 2004).

4. T. J. Richards, D. Ortiz-Barrientos, Immigrant inviability produces a strong barrier to gene flow between parapatric ecotypes of *Senecio lautus*. Evolution 70, 1239–1248 (2016).

5. C. Pinho, J. Hey, Divergence with Gene Flow: Models and Data. Annu. Rev. Ecol. Evol. Syst. 41, 215–230 (2010).

6. E. Baack, M. C. Melo, L. H. Rieseberg, D. Ortiz-Barrientos, The origins of reproductive isolation in plants. New Phytol. 207, 968–984 (2015).

7. M. Kirkpatrick, V. Ravigné, Speciation by natural and sexual selection: models and experiments. Am. Nat. 159 Suppl 3, S22–35 (2002).

8. D. I. Bolnick, B. M. Fitzpatrick, Sympatric Speciation: Models and Empirical Evidence. Annu. Rev. Ecol. Evol. Syst. 38, 459–487 (2007).

9. H. Hipperson, L. T. Dunning, W. J. Baker, Ecological speciation in sympatric palms: 2. Pre-and post-zygotic isolation. Journal of Evolutionary Biology 29, 2143–2156 (2016).

10. H. D. Rundle, P. Nosil, Ecological speciation. Ecol. Lett. 8, 336–352 (2005).

11. K. Christie, S. Y. Strauss, Reproductive isolation and the maintenance of species boundaries in two serpentine endemic jewelflowers. Evolution 73, 1375–1391 (2019).

12. C. Epling, Actual and potential gene flow in natural populations. Am. Nat. 81, 104–113 (1947).

13. R. G. Harrison, The language of speciation. Evolution 66, 3643–3657 (2012).

14. J. R. Ehleringer, C. Clark, “Evolution and adaptation in Encelia (Asteraceae)” in Plant Evolutionary Biology, L. D. Gottlieb, S. K. Jain, Eds. (Springer Netherlands, 1988), pp. 221–248.

15. S. D. Fehlberg, T. A. Ranker, Phylogeny and Biogeography of *Encelia* (Asteraceae) in the Sonoran and Peninsular Deserts Based on Multiple DNA Sequences. Syst. Bot. 32, 692–699 (2007).

16. D. W. Kyhos, C. Clark, W. C. Thompson, The Hybrid Nature of *Encelia laciniata* (Compositae: Heliantheae) and Control of Population Composition by Post-Dispersal Selection. Syst. Bot. 6, 399–411 (1981).

17. D. W. Kyhos, Evidence of different adaptations of flower color variants of *Encelia farinosa* (Compositae). Madroño 21, 49–61 (1971).

18. C. Clark, Phylogeny and Adaptation in the *Encelia* Alliance (Asteraceae: Helliantheae). Aliso: A Journal of Systematic and Evolutionary Botany 17, 89–98 (1998).

19. N. H. Barton, G. M. Hewitt, Analysis of hybrid zones. Annu. Rev. Ecol. Syst. 16, 113–148 (1985).

20. J. C. Beatley, Phenological Events and Their Environmental Triggers in Mojave Desert Ecosystems. Ecology 55, 856–863 (1974).

21. G. Vasey, J. N. Rose, List of plants collected by Dr. Edward Palmer in Lower California and western Mexico in 1890. Reports on collections, and miscellaneous papers (1890).

22. J. G. Kingsolver, et al., The strength of phenotypic selection in natural populations. Am. Nat. 157, 245–261 (2001).

23. J. Hereford, A quantitative survey of local adaptation and fitness trade-offs. Am. Nat. 173, 579–588 (2009).

24. T. J. Thurman, R. D. H. Barrett, The genetic consequences of selection in natural populations. Mol. Ecol. 25, 1429–1448 (2016).

25. D. B. Lowry, J. L. Modliszewski, K. M. Wright, C. A. Wu, J. H. Willis, The strength and genetic basis of reproductive isolating barriers in flowering plants. Philos. Trans. R. Soc. Lond. B Biol. Sci. 363, 3009–3021 (2008).

26. D. Hillel, Introduction to Environmental Soil Physics (Elsevier, 2003).

27. I. Noy-Meir, Desert ecosystems: environment and producers. Annu. Rev. Ecol. Syst. 4, 25–51 (1973).

28. R. L. Andrew, K. L. Ostevik, D. P. Ebert, L. H. Rieseberg, Adaptation with gene flow across the landscape in a dune sunflower. Mol. Ecol. 21, 2078–2091 (2012).

29. H. Lambers, F. Stuart Chapin III, T. L. Pons, Plant Physiological Ecology (Springer Science & Business Media, 2008).

30. P. J. Kramer, J. S. Boyer, Water Relations of Plants and Soils (Academic Press, 1995).

31. R. Bhaskar, A. Valiente-Banuet, D. D. Ackerly, Evolution of hydraulic traits in closely related species pairs from Mediterranean and nonMediterranean environments of North America. New Phytol. 176, 718–726 (2007).

32. J. M. M. De Nava, D. S. Gorsline, G. A. Goodfriend, V. K. Vlasov, R. Cruz-Orozco, Evidence of Holocene climatic changes from aeolian deposits in Baja California Sur, Mexico. Quat. Int. 56, 141–154 (1999).

33. D. Koračin, C. E. Dorman, E. P. Dever, Coastal Perturbations of Marine-Layer Winds, Wind Stress, and Wind Stress Curl along California and Baja California in June 1999. J. Phys. Oceanogr. 34, 1152–1173 (2004).

34. R. A. Bagnold, The Physics of Blown Sand and Desert Dunes (Courier Corporation, 2012).

35. J. F. Kok, E. J. R. Parteli, T. I. Michaels, D. B. Karam, The physics of wind-blown sand and dust. Rep. Prog. Phys. 75, 106901 (2012).

36. S. Vogel, Leaves in the lowest and highest winds: temperature, force and shape. New Phytol. 183, 13–26 (2009).

37. M. A. Maun, Adaptations of plants to burial in coastal sand dunes. Can. J. Bot. 76, 713–738 (1998).

38. F. Roda, et al., Convergence and divergence during the adaptation to similar environments by an Australian groundsel. Evolution 67, 2515–2529 (2013).

39. B. W. Wells, I. V. Shunk, Salt Spray: An Important Factor in Coastal Ecology. Bull. Torrey Bot. Club 65, 485–492 (1938).

40. M. C. Shannon, “Adaptation of Plants to Salinity” in Advances in Agronomy, D. L. Sparks, Ed. (Academic Press, 1997), pp. 75–120.

41. T. J. Flowers, T. D. Colmer, Salinity tolerance in halophytes. New Phytol. 179, 945–963 (2008).

42. P. Proksch, M. Proksch, W. Weck, E. Rodriguez, Localization of chromenes and benzofurans in the genus *Encelia* (Asteraceae). Z. Naturforsch. C 40, 301–304 (1985).

43. P. A. Nakata, Advances in our understanding of calcium oxalate crystal formation and function in plants. Plant Sci. 164, 901–909 (2003).

44. P. D. Coley, J. P. Bryant, F. S. Chapin 3rd, Resource availability and plant antiherbivore defense. Science 230, 895–899 (1985).

45. R. L. Woodman, G. W. Fernandes, Differential Mechanical Defense: Herbivory, Evapotranspiration, and Leaf-Hairs. Oikos 60, 11–19 (1991).

46. M. A. Sorensen, J. A. Bethke, R. A. Redak, Potential host plants of *Trirhabda geminata* (Coleoptera: Chrysomelidae): impacts on survival, development, and feeding. Environ. Entomol. 39, 159–163 (2010).

47. L. Benson, E. A. Phillips, P. A. Wilder, Evolutionary sorting of characters in a hybrid swarm. I: Direction of slope. Am. J. Bot. 54, 1017–1026 (1967).

48. R. I. Milne, S. Terzioglu, R. J. Abbott, A hybrid zone dominated by fertile F1s: maintenance of species barriers in *Rhododendron*. Mol. Ecol. 12, 2719–2729 (2003).

49. C. Christe, et al., Selection against recombinant hybrids maintains reproductive isolation in hybridizing *Populus* species despite F1 fertility and recurrent gene flow. Mol. Ecol. 25, 2482–2498 (2016).

50. M. C. Melo, A. Grealy, B. Brittain, G. M. Walter, D. Ortiz-Barrientos, Strong extrinsic reproductive isolation between parapatric populations of an Australian groundsel. New Phytol. 203, 323–334 (2014).

51. J. M. Sobel, G. F. Chen, L. R. Watt, D. W. Schemske, The biology of speciation. Evolution 64, 295–315 (2010).

52. W. R. Rice, E. E. Hostert, Laboratory experiments on speciation: what have we learned in 40 years? Evolution 47, 1637–1653 (1993).

53. M. R. Servedio, M. A. F. Noor, The Role of Reinforcement in Speciation: Theory and Data. Annu. Rev. Ecol. Evol. Syst. 34, 339–364 (2003).

54. D. A. Levin, Reinforcement of Reproductive Isolation: Plants Versus Animals. Am. Nat. 104, 571–581 (1970).

55. R. Hopkins, M. D. Rausher, Pollinator-mediated selection on flower color allele drives reinforcement. Science 335, 1090–1092 (2012).

56. S. Gavrilets, Fitness Landscapes and the Origin of Species (Princeton University Press, 2004).

57. O. Seehausen, Conservation: losing biodiversity by reverse speciation. Curr. Biol. 16, R334–7 (2006).

58. D. K. Jacobs, T. A. Haney, K. D. Louie, Genes, diversity, and geological process on the Pacific coast. Annu. Rev. Earth Planet. Sci. 32, 601–652 (2004).

59. D. R. Zerbino, E. Birney, Velvet: algorithms for de novo short read assembly using de Bruijn graphs. Genome Res. 18, 821–829 (2008).

60. T. Rognes, T. Flouri, B. Nichols, C. Quince, F. Mahé, VSEARCH: a versatile open source tool for metagenomics. PeerJ 4, e2584 (2016).

61. H. Li, Aligning sequence reads, clone sequences and assembly contigs with BWA-MEM. arXiv [q-bio.GN] (2013).

62. T. S. Korneliussen, A. Albrechtsen, R. Nielsen, ANGSD: Analysis of Next Generation Sequencing Data. BMC Bioinformatics 15, 356 (2014).

63. L. Skotte, T. S. Korneliussen, A. Albrechtsen, Estimating individual admixture proportions from next generation sequencing data. Genetics 195, 693–702 (2013).

64. E. P. Derryberry, G. E. Derryberry, J. M. Maley, R. T. Brumfield, HZAR: hybrid zone analysis using an R software package. Mol. Ecol. Resour. 14, 652–663 (2014).

65. C. R. Brodersen, A. B. Roddy, New frontiers in the three-dimensional visualization of plant structure and function. Am. J. Bot. 103, 184–188 (2016).

66. C. R. Brodersen, Visualizing wood anatomy in three dimensions with high-resolution X-ray micro-tomography (μCT) – a review. IAWA Journal 34, 408–424 (2013).

