## Supplementary Information for "Natural selection maintains species despite widespread hybridization in the desert shrub *Encelia*"

### SUPPLEMENTAL METHODS

#### Site Characteristics

All sampling, field measurements, and experiments for *E. palmeri* - *E. ventorum* were carried out within the Reserva de la Biosfera El Vizcaíno, at a site 1 km south of the settlement of San Roque, 5 km north of the town of Bahía Asunción, on the Pacific Coast of Baja California Sur, México (Figure 1; N 27.17675, W 114.37088). All sampling for *E. asperifolia* - *E. ventorum* were carried out on a site on the Pacific Coast of Baja California Sur, México (Figure 1; N 28.887, W 114.4285). Both sites are characteristic of the sand dune-desert ecotones found throughout the Baja California Peninsula.

The region is characterized by a coastal desert climate, with low annual precipitation, strong coast-inland summer temperature gradients, and frequent summer fog (1–3). Rainfall averages less than 80 mm per year (WorldClim 30-second precipitation maps, http://www.worldclim.org/) with the majority of precipitation falling in the winter as the remnants of frontal systems originating in the north Pacific Ocean, although localized convective storms can contribute significant amounts of summer rain (1, 2).

Desert habitats in the study areas are composed of recently deposited mid-Miocene marine sediments eroded from nearby terraces uplifted during the Pleistocene, with frequent outcrops of Pliocene and Pleistocene volcanic deposits (3–5). Sand dune habitats are created wherever onshore winds are able to transport beach sand inland (6, 7). The consistent afternoon onshore winds at the study site (Figure S11) are characteristic of most of the Pacific Coast of the Baja California Peninsula (2, 7, 8). The predictability of the wind directionality creates stable, linear ecotones between dune and desert habitats with dunes moving parallel to the ecotone (Figure S1). Ecotones are thus defined by the tangent of the prevailing wind direction to the curvature of the beach, which is the source of the dune sand (9, 10).

**Hybrid zone analysis**

To analyze patterns of hybridization and introgression at the *E. palmeri* - *E. ventorum* and *E. asperifolia* - *E. ventorum* hybrid zones, we analyzed phenotypic and genetic data from transects across each hybrid zone.

To sample the hybrid zones, we collected leaves for genetic and phenotypic analysis from adult plants in each habitat and in the ecotone. For each individual, we sampled multiple adult leaves and recorded both latitude & longitude and habitat type. We followed a transect through the hybrid zone, sampling perpendicular to the transect in the ecotone due to the narrowness of the transition. In total, we sampled 112 individuals for the *E. palmeri* - *E. ventorum* hybrid zone and 91 individuals for the *E. asperifolia* - *E. ventorum* hybrid zone.

To characterize genetic patterns at the hybrid zones, we collected double-digest restriction aided (ddRAD) data from each individual. First, we used Qiagen DNeasy Plant Kit (cat no: 69181) to extract DNA, following the manufacturer's instructions. Then, we contracted RTL Genomics (Lubbock, TX, USA) to prepare ddRAD libraries for each individual. DNA samples were cut with the restriction enzymes *PstI* and *MspI*, size-selected from 250 – 700 bp, and then doubly-barcoded to produce uniquely-barcoded libraries. We combined equimolar amounts of barcoded libraries to produce a single pool per hybrid zone. These pools were then sequenced across one lane of 100PE reads on the

Illumina HiSeq 4000 platform at the Vincent Coates Genomics Sequencing Laboratory (Berkeley, CA).

To analyze these genetic data, we first demultiplexed and trimmed all reads of adaptor using Trimmomatic v36 (11). Per individual, we then merged all overlapping paired-end reads using PEAR v0.9.6 (12) and assembled these reads using Velvet v1.2.10 (13). We then generated a pseudo-reference genome (PRG) per hybrid zone. To do so, we identified all contigs that were assembled across >50% of sampled individuals, using a 95% similarity search in vsearch v2.4.3 (14). We then mapped reads from each individual to this PRG using bwa v0.7.17 (15) and called variants across all individuals in a given hybrid zone using ANGSD v0.923 (16). To determine patterns of admixture and introgression across individuals, we first filtered these variants to retain variants with <50% missing data and then used NGSadmix v32 to calculate ancestry proportions for each individual (17). We used HZAR to estimate genetic clines for each hybrid zone (18). To do so, we lumped individuals into 10-meter bands along the transect, calculating the average ancestry for each band (our “population”). We then fit a sigmoidal cline with fixed tails ($p = 0$ & $p = 1$) using the MCMC method implemented in HZAR.

To collect and analyze phenotypic data from the hybrid zones, we measured leaf area and shape by photographing leaves with a scale bar using a digital camera. Hybrids are phenotypically intermediate between parental taxa, which is evident from a few diagnostic leaf traits (Fig. 1). The ‘analyze particles’ function and ‘solidity > analyze particles’ functions in Image-J (U.S. National Institutes of Health, http://imagej.nih.gov/ij/), were used to measure leaf area and shape, respectively.

After conducting a scaled and centered PCA on all leaves from a given hybrid zone, we averaged PC scores per individual across all leaves measured for that individual. PC1 captured 52% of the variation in the *E. asperifolia* - *E. ventorum* hybrid zone and 45% of the variation in the *E. palmeri* - *E. ventorum* hybrid zone. In both cases, PC1 largely reflected leaf shape.

**Estimates of current hybridization**

Hybridization rates were estimated in 2016 by collecting fruits (cypselae) from phenotypically pure individuals of *E. palmeri* - *E. ventorum* growing within 200 meters of the center of the hybrid zone and assaying their progeny for traits indicative of interspecific hybridization. A large number of cypselae were collected from *E. palmeri* and *E. ventorum* and spread evenly onto 11 × 21 × 3 inch greenhouse flats filled with potting soil in 2016. Flats were placed in a greenhouse at the University of California, Riverside, in 2016. Flats were watered daily for approximately two months until all germination ceased. Seedlings that possessed leaf traits diagnostic for both parental taxa (pubescence and dentition) and that visually scored between -1 and 1 on the hybrid index (Figure 1c) were recorded as hybrid. Of the planted seeds, 175 plants from *E. palmeri* mothers germinated of which 55 (31.4%) exhibited phenotypes indicative of hybridization or backcrossing. For *E. ventorum* mothers, 409 seedlings germinated of which 23 (5.6%) exhibited phenotypes indicative of F1 hybridization or backcrossing.

**Environmental characterization of the *E. palmeri* - *E. ventorum* hybrid zone**

Microclimates in dune and desert habitats were measured using paired weather stations and soil moisture arrays. Weather stations were located at the San Roque experimental site 250 meters apart and on opposite sides of the ecotone, among the experimental transplants (Figure S1). Weather stations were constructed of tubular steel tripods with cross bars mounted one meter off the ground to which the various sensors were attached. Relative humidity, temperature, wind speed, and leaf wetness were measured in dune and desert habitats, while photosynthetic photon flux density and wind direction were measured at the dune location only. Weather stations used Li-Cor and ONSET probes coupled to Hobo data loggers (https://www.onsetcomp.com/) and were programmed to log every 15 minutes. Soil moisture stations used ECH2O soil moisture probes coupled to ONSET Hobo data loggers and recorded soil moisture at -2.0 m, -1.0 m, -0.5 m, and -0.2 meters depth at 30 minute intervals (Figure S3).

*Soils* Soil texture was measured by establishing transects among the experimental plants at the San Roque experimental site. Soil was sampled every 10 meters along seven replicate 200 meter long transects centered on and perpendicular to the ecotone. At each sampling station approximately one liter of soil was collected from between 10 and 20 centimeters below the soil surface. All large organic debris was removed by hand, and the soil dried for two weeks in a low humidity environment until registering a constant mass on a ten kilogram Mettler digital balance with 0.01 gram resolution. Soil was then sifted through U.S. Standard #35 sieve and the throughfall passed through a #60 sieve. The retained fraction thus contained particles from between 0.50 and 0.25 mm in diameter, which is classified as "fine sand" (9). The fine sand fraction was weighed, and the proportion of fine sand calculated as a fraction of the total dry mass. The results were plotted as a function of distance and the arithmetic mean of all measurements for a given distance shown with error bars representing ± one standard error (Figure 3a).

*Community composition* The composition of all other non-*Encelia* vascular perennial plant species at the San Roque experimental site was quantified by recording the abundance of taxa encountered within seven 10 × 200 meter belt transects established among the experimental plants, with data binned at five meter intervals. Non-metric multidimensional scaling (NMDS) was performed on the resulting community data matrix using the "vegdist" function in the "vegan" package, and the "isoMDS" function in the "mass" package in R. Scores from the first NMDS axis for each five meter increment were then averaged across all transects and plotted as a function of distance with error bars representing ± one standard error (Figure 3d). A total of eleven perennial non-*Encelia* vascular plant species were encountered during community sampling of both dune and desert habitats (Table S4), with a near-complete turnover of species composition over a distance of 20-40 meters.

**Reciprocal Transplant**

A reciprocal transplant field experiment was conducted during the winter and spring of 2010 and 2011 at the San Roque experimental site. Parental and hybrid phenotypes grown from seed were planted into dune, desert, and ecotone habitats, and growth, survival, and reproduction measured over the course of one growing season. To obtain parental plants, we collected fertile cypselae throughout spring 2010 from phenotypically pure parental *E. palmeri* and *E. ventorum* individuals, as well as individuals that were phenotypically hybrid. Cypselae were germinated outside in greenhouse flats filled with washed sand and transplanted to the field when they were between 5 and 10 cm tall or had at least 5 true leaves. Hybrids for the reciprocal transplant experiment were obtained from hybrid mothers, and thus exhibited a range of phenotypes indicative of recombination in F2 or later generation hybrids or backcrosses (Figure S12). Parental taxa largely bred true and only progeny exhibiting pure parental phenotypes (i.e. those resulting from intraspecific matings) were used in the experiment. Hybrids were easily distinguishable in the field, with hybrids identified visually by possessing combinations of pubescence and leaf dentation, traits diagnostic for *E. palmeri* and *E. ventorum*, respectively.

A total of 276 plants were reciprocally planted into dune, desert, and ecotone habitats from September to November 2010. The total number of plants in each habitat was 105 in the desert (38 *E. palmeri*, 23 hybrids, 44 *E. ventorum*), 43 in the ecotone (12 *E. palmeri*, 20 hybrids, 11 *E. ventorum*), and 123 in the dunes (54 *E. palmeri*, 20 hybrids, 49 *E. ventorum*). Sample sizes were dictated by the availability of experimental plants, particularly hybrids, and were distributed disproportionately in dune and desert habitats in order to concentrate statistical power at the ends of the hypothesized selective gradients. Planting order was randomized by habitat and phenotype, with individuals that were planted later due to later emergence from the germination flats watered correspondingly longer to standardize the total time spent under irrigation (linear regression of total ln(1+$x$) transformed biomass versus planting date was not statistically significant; $R^2$=0.013, $p$=0.355). Cages were placed around experimental plants to deter disturbance from coyotes. Cages were 0.5 meters high by 0.5 meters in diameter and were made of 1 × 2 inch welded wire mesh (Figures S7, S9). Cages did not substantially affect solar insolation or wind speed, and rabbits and hares are the only known herbivores at the experimental site that may have been prevented access to the experimental plants. All other known herbivores are smaller than the mesh size or could enter through the open top. Only *E. ventorum* individuals in the water addition treatment in the desert were capable of growing large enough to expand outside their cages (see *Water Addition*, below).

Supplemental water was applied to simulate conditions that exist during a typical recruitment year, which occurs approximately every five or ten years. Watering consisted of ten liters of water per plant every other day, applied via backpack-mounted spray wand at a rate of approximately two liters per minute, applied directly over the plant with a spray circumference of approximately 0.5 meters. The watering treatment elevated local soil moisture for approximately 24-36 hours, but thereafter dissipated into the surrounding soil matrix based on test measurements with soil moisture sensors. Watering was terminated beginning in December 2010 and all plants allowed to grow under ambient conditions for four additional months, except for plants in the water addition experiment (see *Water Addition*, below) that received water as described above for the remainder of the experiment. From March 11-16, 2011 the experiment was terminated and all experimental plants were harvested. Stem basal diameter was measured with digital calipers as the average of two perpendicular measurements, one of which was the widest point on the stem, made one centimeter above the soil surface. Plant height and two perpendicular canopy diameters, one of which was the widest possible canopy diameter, were measured with a meter stick. Aboveground and belowground biomass were harvested by excavating plants to a depth of one meter with a trowel and sifting excavated soil through a sieve, removing roots with tweezers. When possible, mid-day shoot water potentials were measured on healthy, 10 cm long shoots with a Scholander-type pressure bomb using methods described in *Hydraulic Physiology*, below. Leaf color, thickness, area, and shape were measured as described below (see *Hybrid Index*) for all plants with healthy foliage. For all plant-level traits, values used in statistical analysis are the average of four mature leaves per plant where possible (range 1-11 leaves) except water potential measurements that were made on one shoot per plant where possible.

Growth was measured as total above plus belowground biomass at the end of the experiment, or the total biomass of the plant at the time of its death. Biomass was ln(1+*x*) transformed in order to improve normality (see *Statistical Analysis*, below), grouped by maternal phenotype, and plotted with standard errors (Figure 4a). Survival was calculated as percent of the total number of individuals planted for each phenotype-habitat combination during the course of the experiment that were alive at the end of the experiment. Reproductive output was measured but only one plant from the water addition experiment flowered during the course of the experiment, so reproductive output was not analyzed. Binomial confidence intervals were calculated for the survival data using the "binom.confint" command in the "binom" library, implemented in R (Figure 4b). Growth and survival were additionally combined into a composite fitness measure by multiplying a binary (0, 1) survival score with the ln(1+*x*) transformed biomass, and taking the arithmetic mean within each phenotype-habitat combination. This is a reasonable fitness approximation because previous research has shown that the single most important predictor of seed production is total plant size (19). In support of this approximation, the one individual to flower during the experiment was the largest plant in the experiment, an *E. ventorum* individual growing in the desert under elevated water availability. The relative fitness data were then relativized within each habitat by setting the phenotype with the highest fitness equal to one and scaling the other phenotypes proportionately (Figure 4c). Negative selection coefficients discussed in the text are calculated as one minus the relative fitness (20). Nonparametric 95th percentile confidence intervals for relative fitness were simulated with a bootstrap procedure using the "boot.ci" function of the "boot" package in R, using 1000 iterations.

*Hybrid Index* In order to more precisely quantify the degree of morphological variability, we measured leaf shape, color, thickness, and area for all plants used in the experiment. Leaf thickness was measured with digital calipers (Mitutoyo Model 700-126, 0.1 mm resolution) on three healthy leaves per plant by taking the arithmetic mean of two measurements, one at the leaf tip and one in the middle of the lamina, avoiding any major veins. Leaf size was measured by photographing leaves with a scale bar using a digital camera. Images were then converted to 8-bit grayscale in Image-J (U.S. National Institutes of Health, http://imagej.nih.gov/ij/), thresholded, and area of the leaf lamina measured using the "analyze particles" function. Leaf color was measured by adjusting the 8-bit grayscale images for brightness by using "brightness adjust" toolbar in Adobe Photoshop CS6 to bring the black and white reference to a constant brightness.

Leaf lamina brightness was then read as the arithmetic mean on a 0-255 scale of all pixels contained within the largest circumscribed rectangle possible within the center of the leaf, excluding from analysis any holes or necrotic or discolored tissue. Leaf shape was characterized by selecting "solidity" in the "analyze particles" function in Image-J, which places the smallest convex hull possible around the object and then calculates the proportion from 0 to 1 inclusive of the area of the hull occupied by leaf lamina. Using this metric, *E. ventorum* exhibits low solidity and *E. palmeri* exhibits high solidity. All variables were tested for normality using Shapiro tests and for homogeneity of variances using Fligner tests implemented in R (R Core Development Team, version 2.15.2), and all were found to be within reasonable limits (data not shown), thus untransformed data was used for all subsequent analyses.

Principal components analysis (PCA) was then performed on the untransformed trait data matrix using the "princomp" function in R. Plotting the first two principal component axes revealed that the first axis was strongly associated with leaf shape, color, and thickness, while the second axis was strongly associated with leaf size (Figure S12). Taxa visually identified as pure parentals exhibited discrete phenotypes that separated along axis one, with hybrids (identified as possessing combinations of leaf dentition and leaf pubescence) spanning the range of phenotypic variation observed between parental taxa.

*Statistical Analysis* To statistically test for the effects of habitat and phenotype on biomass we first tested the assumptions required for parametric analysis by plotting the raw biomass data as well as the ln(1+$x$) transformed biomass data in histograms and conducting Shapiro tests of normality in R. In the desert habitat each phenotype had a biomass distribution that was significantly different than normal for the untransformed data, and not significantly different than normal for the transformed data (results not shown). For all phenotypes in the ecotone, all of the biomass distributions were significantly different than normal even after transformation. The failure of transformations to correct for non-normality is likely caused by the low statistical power in the ecotone compared to dune or desert habitats due to the low sample size in that habitat. Despite this, the deviation from normality was ameliorated substantially in all cases by transformation, and visual inspection of the data indicated that most parametric tests would be robust against deviations of this magnitude. In the dune habitat *E. palmeri* was the only phenotype that exhibited normally distributed biomass despite a low number of individuals surviving to the end of the experiment in the dunes. Shapiro tests and visual inspection of the transformed biomass data from the dune habitat revealed that deviations from normality among all phenotypes were not likely to seriously violate the assumptions of parametric tests and so we proceeded with parametric analysis of the transformed biomass data.

To test for effects of habitat and phenotype on total biomass, based on the above analyses we chose to analyze all the data simultaneously in a general linear model with Gaussian distributed variance and link functions, using phenotype and habitat as fixed independent variables and ln(1+biomass) as the dependent variable. We performed backwards stepwise model selection comparing Akaike Information Criterion (AIC) scores and using analysis of deviance (AOD) to test for differences between models (Table S1). Not all combinations of interactions and variables were tested due to the large number of models this would generate and concerns about multiple comparisons, thus we selected *a priori* models with and without the specific interactions and compared these to saturated models and models without interactions (Table S1). Models were constructed and AIC scores calculated using the "glm" command in R. Analysis of deviance was then performed on the two models with the lowest AIC scores by using the "anova" command in R. For growth, the model including an interaction between phenotype and habitat was a significantly better fit than a model without the interaction (Table S1). Analysis of deviance was performed to compare the fit of the two models, which detected a significantly worse fit when omitting the habitat-by-phenotype interaction ($p$ = 0.0042, $F$ = 4.0704, df = 4). Due to the significant interaction we did not pursue further testing of individual effects.

To statistically test for the effects of habitat and phenotype on survival, we performed backwards stepwise log-linear analysis on three-way contingency tables since our counts of alive versus dead individuals did not meet the requirements of parametric analysis. To assemble the contingency tables we recorded whether each plant was alive or dead at the end of the experiment and coded each observation 0 for dead and 1 for alive and grouped them by habitat (desert, ecotone, and dune) and phenotype (*E. palmeri*, hybrid, and *E. ventorum*) resulting in a 3 × 3 contingency table with the count of plants alive in each cell. We then used the "loglm" command in R and built models in a backwards stepwise fashion to test the importance of habitat-by-phenotype interactions in explaining patterns of survival, with model fit assessed by comparing AIC scores and testing the best-fit models with likelihood ratio (LR) tests (Table S1). We found that the model that included an interaction between predictor variables (the saturated model) fit the data better than any other model (Table S1). Likelihood ratio (LR) tests were then conducted to determine whether differences in model fit were statistically significant. We performed a LR test on the comparison between the saturated model and the model without the interaction (the model with the second-best AIC score) and found that the saturated model was significantly better fit ($p$ < 0.001, Δ= 36.84, df = 4).

To test for the effects of habitat and phenotype on fitness, we first plotted the untransformed absolute fitness data as histograms and conducted Shapiro tests on untransformed as well as ln(1+$x$) transformed data. Shapiro tests detected non-normal distributions for all combinations of phenotype and habitat (data not shown), likely due to overdispersion and an excess of zeroes. The negative binomial distribution was tested against our data using Kolmogorov-Smirov tests, which determined that they were not significantly different (results not shown). Thus, we proceeded with a parametric analysis by constructing a general linear model using a negative binomial variance and link function implemented with the "glm.nb" function in R on ln(1+*x*) transformed data. Comparison of AIC scores and residual deviations showed that the best model included a phenotype-by-habitat interaction (Table S1). A likelihood ratio test also supported that the model without the phenotype-by-habitat interaction was a poorer fit than the model with the interaction ($p = 0.0004$, $\Delta = 20.36$, df = 4). Visual examination of the data also supports this conclusion (Figure 4).

**Water Addition**

The importance of water availability on the relative fitness of *Encelia palmeri* and *E. ventorum* was tested by adding supplemental water to a subset of the plants in the reciprocal transplant field experiment. Five plants of *E. palmeri* and eight plants of *E. ventorum* were randomly chosen in each of dune and desert habitats to continue receiving water beyond the initial two month watering period (see *Reciprocal Transplant*, above) for a total of 3 additional months under irrigation. Hybrids and the ecotone were not included due to a shortage of experimental plants. Water was applied to the experimental plants with a backpack-mounted sprayer as described above. Plants were harvested and assayed for biomass and leaf traits at the same time as the rest of the plants in the reciprocal transplant experiment as described above.

To test the effects of the watering treatment on growth, we plotted ln(1+*x*) transformed biomass for each habitat-by-phenotype combination in histograms and visually determined that a negative binomial distribution would be a better fit to the data than a normal or Poisson distribution due to overdispersion and an excess of zeroes (results not shown). We chose to analyze each habitat individually due to the low statistical power available and our interest in the potential conditional neutrality of some sources of selection. To analyze the ln(1+*x*) transformed biomass data, we constructed a general linear model using a negative binomial link function using the "glm.nb" command in R, first specifying a model with a water × phenotype interaction, and then specifying a model with water and phenotype independently added. Our results indicate that the model with the interaction was a better fit than the model without the interaction, as shown by comparison of AIC scores (Table S1) and a likelihood ratio test ($p$ = 0.0063 , Δ = 7.469, df = 91). This conclusion is supported visually by the absence of fitness differences between *E. palmeri* and *E. ventorum* in the desert habitat in the presence of supplemental water (Figure 5b), while *E. palmeri* performs significantly better than *E. ventorum* in the absence of supplemental water (Figure 5a). For the dune habitat, identical model construction yielded different results. The lowest AIC score belonged to the model without a habitat-by-phenotype interaction, and the model with the interaction had the poorest fit of the three models examined. Despite this, LR tests revealed no significant differences between the models.

### Sources of mortality

Experimental plants were visited two to three times per week during the experiment, providing an opportunity to record instances of mortality due to herbivory and burial by wind-blown sand when the cause was unambiguous. Our estimates of the effects of herbivory and burial likely underestimate the true magnitude of the effects of these factors because a plant had to be completely killed and have clear evidence of the cause of death to be scored. Mortality due to herbivory was scored if the entire plant was destroyed between visits, with evidence of herbivore activity nearby such as tracks or the organism itself, and no other proximate cause identified (e.g. bad weather). Mortality due to burial was scored if the entire plant was completely buried by sand and the leaves visibly wilted and lacking turgor when excavated (Figure S7). Dunes regularly shift in elevation up to 0.3 meters per day during wind storms, with the rate of aggradation or degradation depending greatly on the microsite of the experimental plant. We plotted the percentage of plants for each phenotype in each habitat that were scored as killed by burial or herbivory, with error bars representing binomial 95% confidence intervals as described above (Figure S8).

To analyze patterns of burial in the reciprocal transplant experiment, we performed backwards stepwise log-linear analysis on three-way contingency tables as described above. To test for a habitat-by-phenotype interaction, we constructed models as described above and compared AIC scores and performed LR tests on the two best fitting models (Table S1). For burial, we found that a saturated model without the habitat-by-phenotype interaction best fit the data, although LR testing indicated that there was no difference between this and the model with the second lowest AIC score ($p = 0.165$, $\Delta = 6.498$, df = 4), indicating that habitat and taxon independently contribute to mortality, but evidence of an interaction is weak with these sample sizes. To analyze patterns of herbivory in the reciprocal transplant experiment, we performed backwards stepwise log-linear analysis on three-way contingency tables in an identical fashion as for the analysis of burial. To test for a habitat-by-phenotype interaction, we constructed models as described above and compared AIC scores and performed LR tests on the two best fitting models (Table S1). Comparison of AIC scores indicated the model with the habitat-by-phenotype interaction fit the data better than models without the interaction, and a LR test on the saturated model and the model without interaction found a significant difference between these models ($p = 0.029$, $\Delta = 10.766$, df = 4).

**Hydraulic Physiology**

*Leaf Hydraulic Capacitance* For each phenotype, leaf pressure-volume curves were used to determine leaf hydraulic capacitance, which is the slope of the relationship between water potential and the amount of water released per unit dry mass (21, 22). Leaves were collected at the San Roque field site in the early morning, placed in a humidified plastic bag and returned to the laboratory, and allowed to rehydrate in distilled water for between two and four hours. Rehydration allowed leaves to reach their maximum water potential, and compensates for differences in water availability between habitats. After removing leaves from the distilled water we manually dried their exteriors with paper towels and immediately measured their fresh, fully hydrated mass. Leaves were then allowed to desiccate to different degrees in order to induce declines in water potential and water content. Individual leaves from this desiccating population were measured for their partially hydrated mass and immediately placed in stainless steel chambers to measure their water potential using thermocouple psychrometers (Model 83-3V, Merrill Specialty Equipment, Logan, UT, USA; (23)). These chambers were sealed, triple-bagged, and placed in a circulating water bath set to 25 °C for between four and six hours or until sequential voltage measurements had stabilized.

Thermocouple psychrometers were connected to an AM16/32B multiplexer interfaced with a CR6 datalogger (Campbell Scientific, Logan Utah). Voltage output for each chamber was converted to pressure (MPa) based on sensor-specific NaCl calibration curves following standard methods (24). After water potentials had stabilized, leaves were removed from the psychrometer chambers, immediately re-weighed, and kept on silica until they could be oven-dried. Samples were dried at 65 °C for 72 hours and weighed for their final dry mass. Leaf hydraulic capacitance was calculated as the initial slope of the relationship between water content and water potential. For each leaf, the amount of water released was calculated as the difference between fresh mass and the average of the two masses measured just before and after measurements of water potential. The mass of water was converted to moles of water, divided by the final dry mass of the leaf, and plotted against the leaf water potential (Figure S6). Over-saturation due to rehydration produced a plateau in the amount of water released at very high water potentials, thus we removed these points. Linear regressions were fit through the five points with the highest water potential and points with lower water potentials sequentially added until the addition of a point caused the $R^2$ of the linear regression to decline. This initial slope of the curve is defined as the leaf hydraulic capacitance (22), which was normalized by leaf dry mass.

*Stem Hydraulic Conductance* Whole-stem hydraulic conductance was measured using a vacuum method (25). Healthy 30-50 centimeter long shoots were excised from plants grown from seed in a common garden at the Agricultural Operations Station, Riverside, California, and immediately recut under water. Shoots were taken from three individuals per species, covered with aluminum foil, and returned to the lab. Shoots were kept in water until they were measured, which occurred 1-6 hours after harvest. For measurement, shoots were defoliated under water and the cut surface recut immediately prior to placement in the vacuum chamber. Total leaf area was determined for all removed leaves using ImageJ (see *Hybrid Index*, above). The cut stem was connected with a compression fitting to hard wall tubing that led into a beaker of 0.10 M KCl that was placed on a balance with an accuracy of 0.1 mg (Sartorius, Model CPA225D). The shoot was then placed in a vacuum chamber constructed of clear PVC, and the steady-state flow rate measured as the reduction in mass of the beaker recorded while the stem was under different partial vacuum pressures (60, 50, 40, 30, 20, 10 kPa below ambient air). Whole-stem hydraulic conductance was calculated from the linear regression of flow rate as a function of pressure. This measurement was normalized by total leaf area, giving leaf area adjusted whole-shoot hydraulic conductance (Table S2).

_Osmotic Potential_ In March 2015, we assayed the leaf osmotic potentials of nine *E. ventorum* plants growing along a transect from the ecotone to the foredunes (the closest dunes to the ocean) at the San Roque experimental site, sampling individuals approximately every 30 meters until we reached the last adult *E. ventorum* that occurred nearest the desert habitat. The leaves of *E. palmeri* and hybrids were not used because leaves of these phenotypes were not thick enough to be able to excise non-photosynthetic tissues. At each plant we collected healthy, mature leaves from the ocean-facing side of between five and seven different shoots and kept these leaves in a humidified plastic bag until returning them to the laboratory. Once in the laboratory, we measured leaf area, leaf thickness, leaf color, and leaf solidity on five leaves from each plant or position on the foremost dune plant. In *E. ventorum* leaves, the green, photosynthetic mesophyll occurs as a ring around the outside of leaves. Inside this ring of tissue is a white, non-photosynthetic, parenchymous tissue that increased in thickness with greater ocean exposure (Figure S10). We hypothesized that the increase in thickness of this tissue may be associated with lower osmotic potentials that would drive increased water import. To test this, we measured the osmotic potential of this tissue. Under a saturated atmosphere, we manually excised the green, photosynthetic tissue from 3-5 leaves per plant along the transect, macerated the white parenchymous tissue, and placed this pooled sample into a thermocouple psychrometer chamber. Maceration eliminated effects on water potential due to tension in the xylem and turgor pressure, such that the measured water potential was assumed to be equivalent to the osmotic potential of the tissue. Psychrometers were incubated and measurements made as described above and plotted against relative position of the plant along the dune transect.

*Leaf Succulence* Leaf thickness was also measured as a proxy for leaf succulence. Leaf thickness was characterized by sampling plants at approximately 30 meter intervals in a 350 meter long transect running from the exposed foredune to the ecotone at the San Roque experimental site (n = 45; Figure S10b). We additionally measured leaf thickness at five points around the circumference of a large individual of *E. ventorum* growing on the exposed foredune as another test for the effects of ocean exposure on leaf morphology (n = 30; Figure S10c).

### Supplemental References

1. A. F. Johnson, A Survey of the Strand and Dune Vegetation Along the Pacific and Southern Gulf Coasts of Baja California, Mexico. *J. Biogeogr.* **4**, 83–99 (1977).

2. D. K. Jacobs, T. A. Haney, K. D. Louie, Genes, diversity, and geological process on the Pacific coast. *Annu. Rev. Earth Planet. Sci.* **32**, 601–652 (2004).

3. M. Peinado, J. Delgadillo, J. L. Aguirre, Plant associations of El Vizcaíno biosphere reserve, Baja California Sur, Mexico. *Southwest. Nat.* **50**, 129–149 (2005).

4. A. L. Carreño, J. L. M. Ruiz, Diatom biostratigraphy of Bahia Asunción, Baja Califórnia Sur, México. *Revista mexicana de ciencias geológicas* **11**, 11 (1994).

5. J. Ledesma-Vázquez, R. Hernández-Walls, M. Villatoro-Lacouture, R. Guardado-France, Dynamics of rocky shores: Cretaceous, Pliocene, Pleistocene, and Recent, Baja California peninsula, Mexico. *Can. J. Earth Sci.* **43**, 1229–1235 (2006).

6. M. A. Maun, Adaptations of plants to burial in coastal sand dunes. *Can. J. Bot.* **76**, 713–738 (1998).

7. J. M. M. De Nava, D. S. Gorsline, G. A. Goodfriend, V. K. Vlasov, R. Cruz-Orozco, Evidence of Holocene climatic changes from aeolian deposits in Baja California Sur, Mexico. *Quat. Int.* **56**, 141–154 (1999).

8. D. Koračin, C. E. Dorman, E. P. Dever, Coastal Perturbations of Marine-Layer Winds, Wind Stress, and Wind Stress Curl along California and Baja California in June 1999. *J. Phys. Oceanogr.* **34**, 1152–1173 (2004).

9. R. A. Bagnold, *The Physics of Blown Sand and Desert Dunes* (Courier Corporation, 2012).

10. J. F. Kok, E. J. R. Parteli, T. I. Michaels, D. B. Karam, The physics of wind-blown sand and dust. *Rep. Prog. Phys.* **75**, 106901 (2012).

11. A. M. Bolger, M. Lohse, B. Usadel, Trimmomatic: a flexible trimmer for Illumina sequence data. *Bioinformatics* **30**, 2114–2120 (2014).

12. J. Zhang, K. Kobert, T. Flouri, A. Stamatakis, PEAR: a fast and accurate Illumina Paired-End reAd mergeR. *Bioinformatics* **30**, 614–620 (2014).

13. D. R. Zerbino, E. Birney, Velvet: algorithms for de novo short read assembly using de Bruijn graphs. *Genome Res.* **18**, 821–829 (2008).

14. T. Rognes, T. Flouri, B. Nichols, C. Quince, F. Mahé, VSEARCH: a versatile open source tool for metagenomics. *PeerJ* **4**, e2584 (2016).

15. H. Li, Aligning sequence reads, clone sequences and assembly contigs with

BWA-MEM. *arXiv [q-bio.GN]* (2013).

16. T. S. Korneliussen, A. Albrechtsen, R. Nielsen, ANGSD: Analysis of Next Generation Sequencing Data. *BMC Bioinformatics* **15**, 356 (2014).

17. L. Skotte, T. S. Korneliussen, A. Albrechtsen, Estimating individual admixture proportions from next generation sequencing data. *Genetics* **195**, 693–702 (2013).

18. E. P. Derryberry, G. E. Derryberry, J. M. Maley, R. T. Brumfield, HZAR: hybrid zone analysis using an R software package. *Mol. Ecol. Resour.* **14**, 652–663 (2014).

19. J. R. Ehleringer, C. S. Cook, Characteristics of Encelia species differing in leaf reflectance and transpiration rate under common garden conditions. *Oecologia* **82**, 484–489 (1990).

20. T. J. Thurman, R. D. H. Barrett, The genetic consequences of selection in natural populations. *Mol. Ecol.* **25**, 1429–1448 (2016).

21. H. Lambers, F. Stuart Chapin III, T. L. Pons, *Plant Physiological Ecology* (Springer Science & Business Media, 2008).

22. K. A. McCulloh, D. M. Johnson, F. C. Meinzer, D. R. Woodruff, The dynamic pipeline: hydraulic capacitance and xylem hydraulic safety in four tall conifer species. *Plant Cell Environ.* **37**, 1171–1183 (2014).

23. P. J. Kramer, J. S. Boyer, *Water Relations of Plants and Soils* (Academic Press, 1995).

24. R. W. Brown, D. L. Bartos, A calibration model for screen-caged Peltier thermocouple psychrometers. *Res. Pap. INT-293.* (1982) https:/doi.org/10.2737/int-rp-293.

25. K. J. Kolb, J. S. Sperry, B. B. Lamont, A method for measuring xylem hydraulic conductance and embolism in entire root and shoot systems. *J. Exp. Bot.* **47**, 1805–1810 (1996).

### SUPPLEMENTAL FIGURES

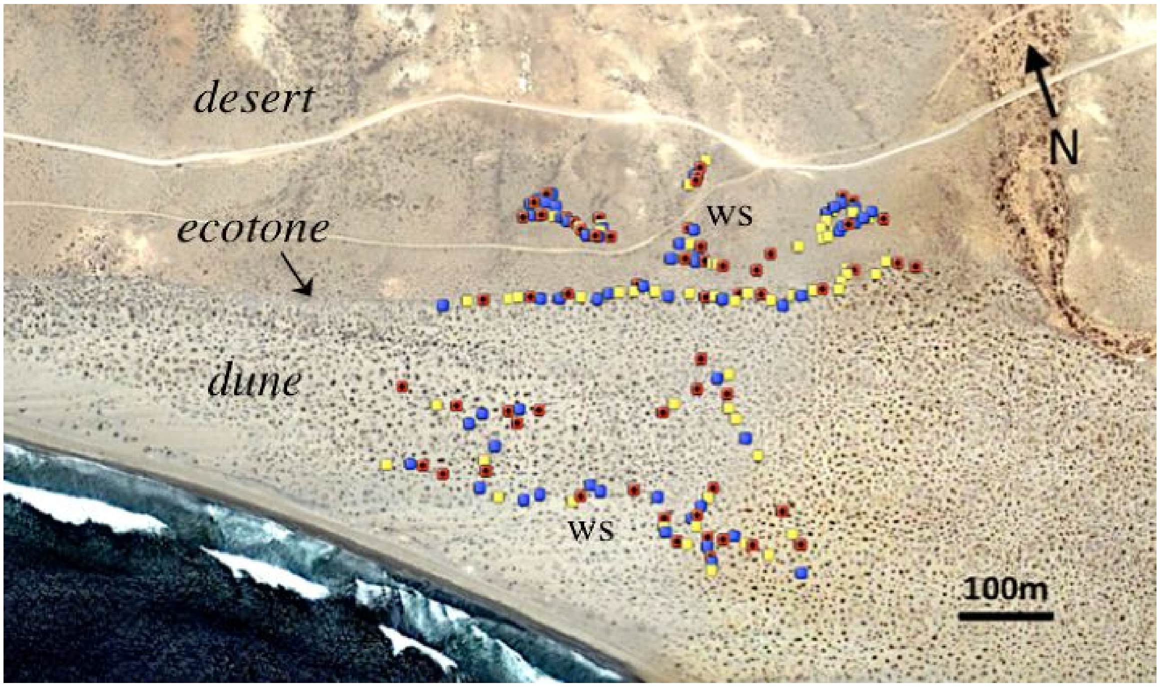

**Figure S1:** Map of the San Roque experimental site with the locations of the dune (top), ecotone (center), and dune (bottom) habitats. Locations of the experimental plants are shown in blue (*E. palmeri*), yellow (hybrids), and red (*E. ventorum*). Locations of weather and soil moisture stations are denoted "WS".

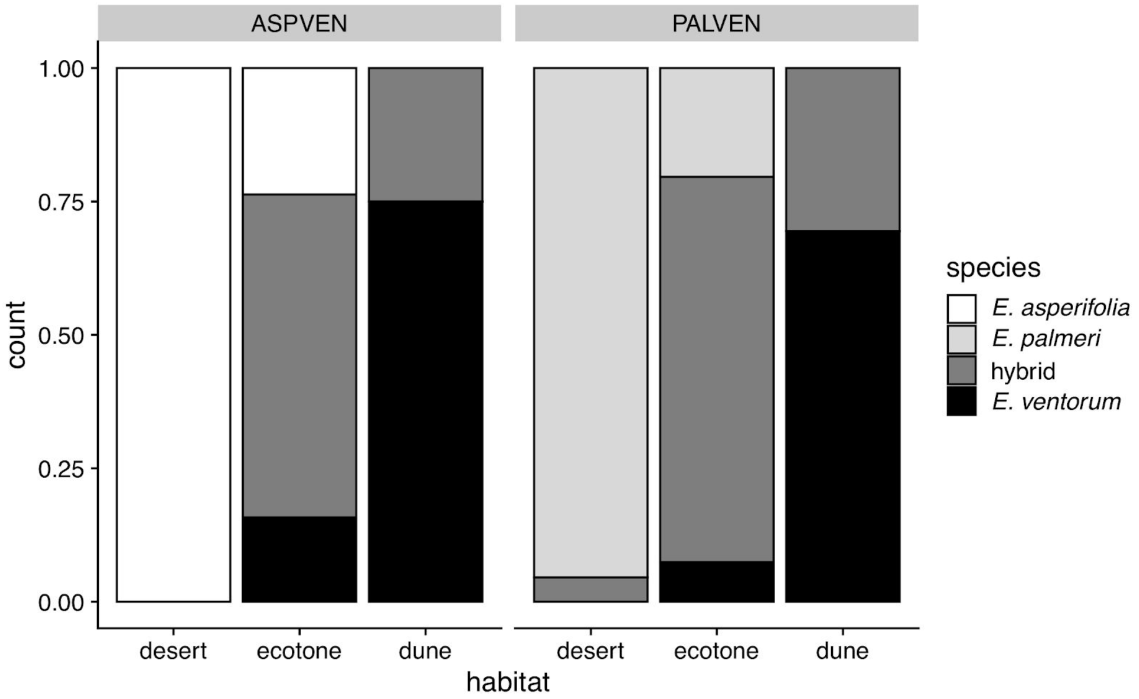

**Figure S2:** Patterns of species composition across habitat types in the *Encelia palmeri* - *E. ventorum* hybrid zone (N = 112; PALVEN) and the *E. asperifolia* - *E. ventorum* hybrid zone (N = 91; ASPVEN). Hybrids were defined as any individual that had >10% genetic ancestry for both parental species. Across both hybrid zones, parental species are most commonly found in their typical habitat, and hybrids are dominant at the ecotone.

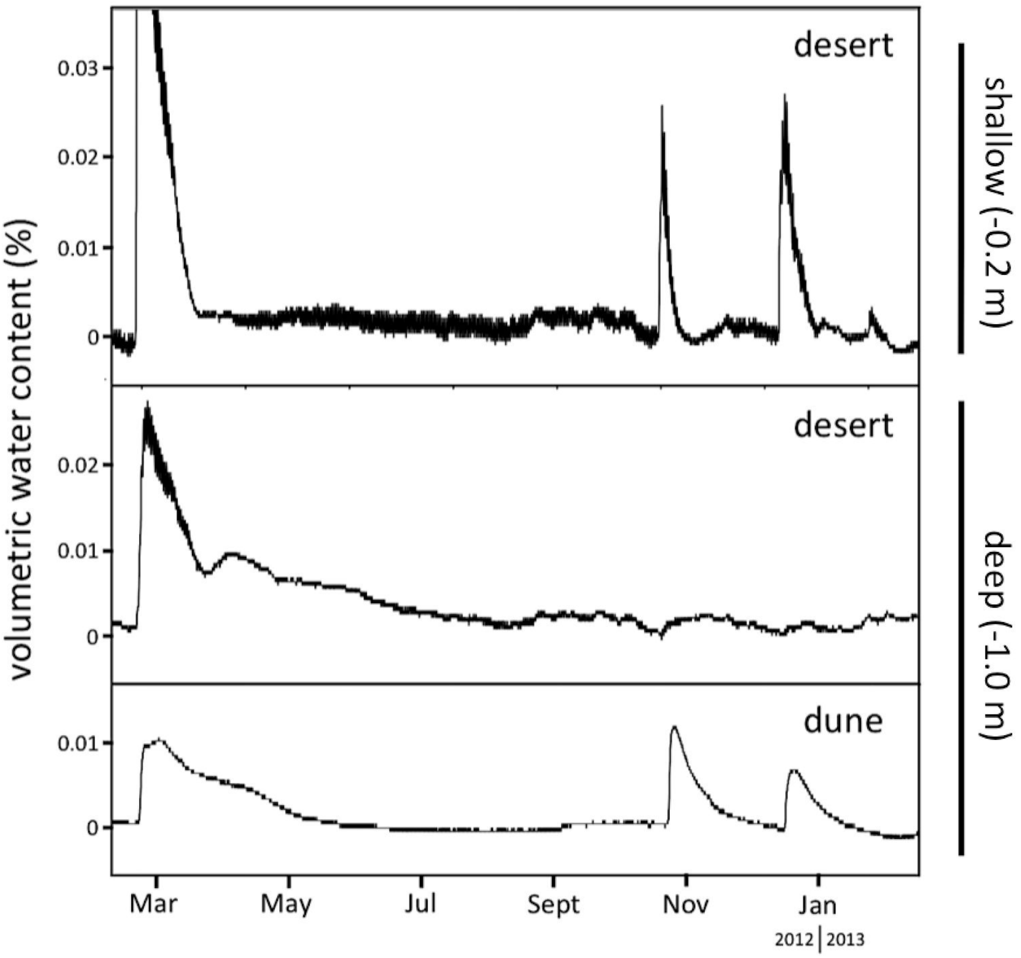

**Figure S3:** Soil water content in dune and desert habitats, at shallow and deep soil horizons, from the San Roque experimental site (*Methods: Site Characteristics*). Two small rain events in October and December 2012 were detected in the shallow desert horizon but were not detected in the deep desert horizon. In contrast, these same two rain events were detected in both the shallow and deep dune horizons, where it persisted for several months. Large but infrequent rain events such as in February 2012 are, however, able to infiltrate into the deep horizon in the desert.

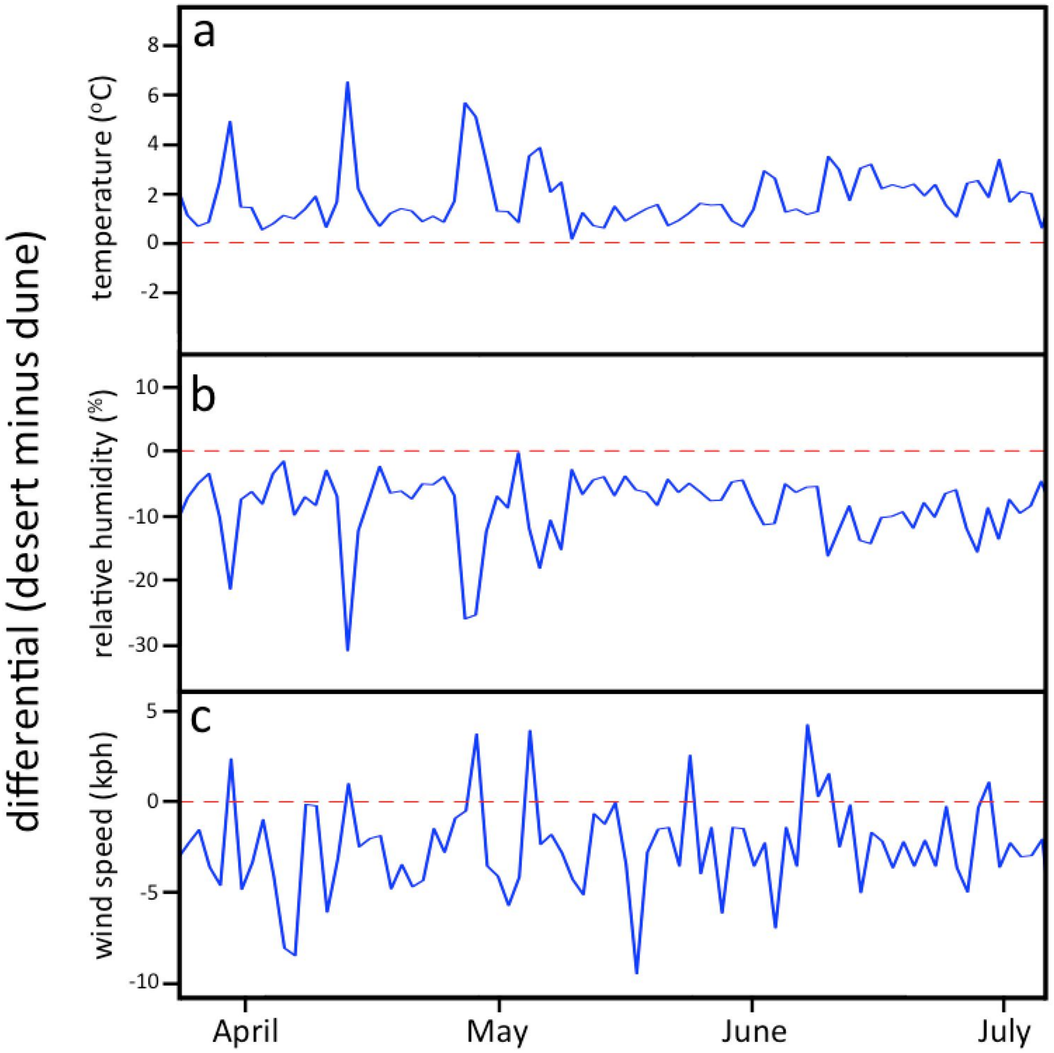

**Figure S4:** Differentials at the San Roque transplant site in spring 2011 between desert and dune habitats for (a) temperature, (b) relative humidity, and (c) wind speed, measured using paired weather stations separated by a distance of 250 meters (Figure S1). Data are restricted to daytime measurements between 15:00 and 16:00, with measurements taken every 15 seconds and averaged for a daily value. The desert habitat is on average 3.20 °C hotter and 9.24% less humid and experiences 2.54 km $h^{-1}$ slower afternoon wind speeds than the dune habitat.

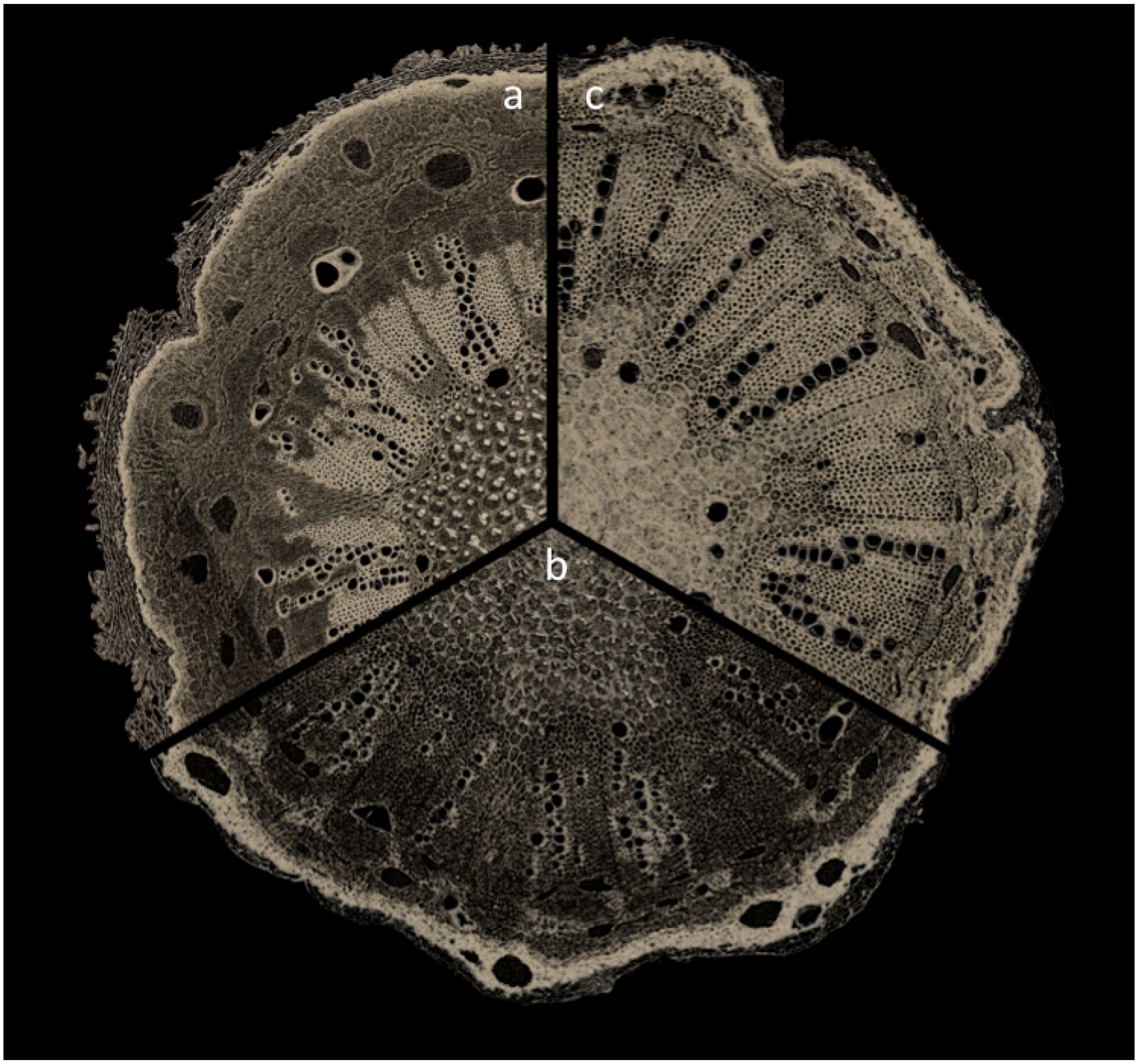

**Figure S5:** Stem cross sections of (a) *E. palmeri*, (b) a hybrid, and (c) *E. ventorum,* illustrating variation in xylem vessel diameter. Stem diameters are between three and five mm.

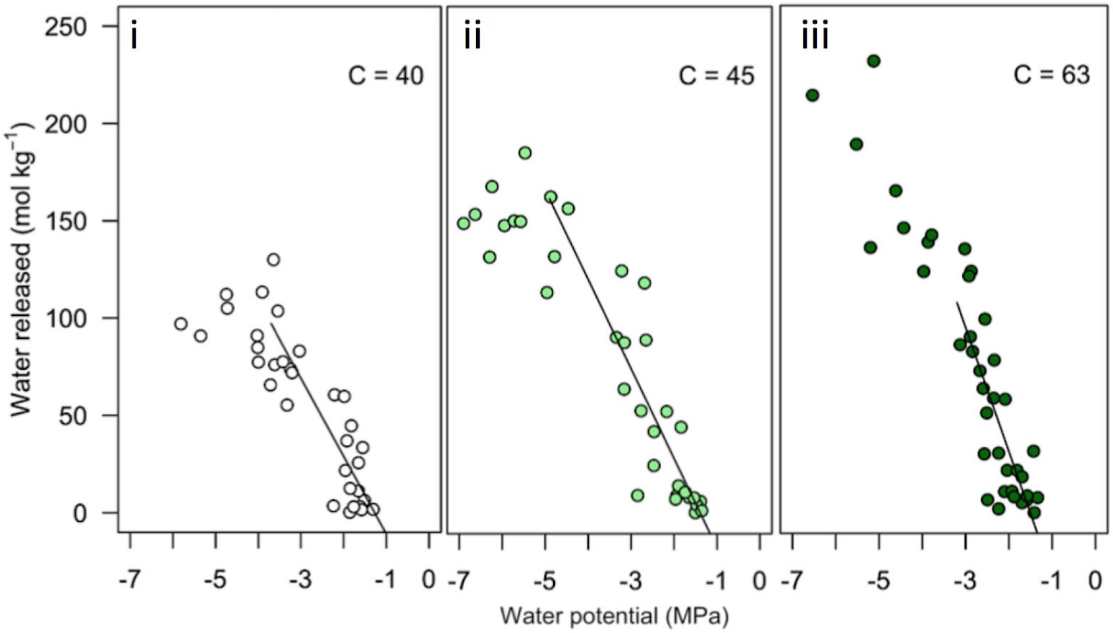

**Figure S6:** Leaf hydraulic capacitance (C), for (i) *Encelia palmeri*, (ii) natural hybrids, and (iii) *E. ventorum*, measured as the slope of the best-fit line relating the decrease in water potential for a given reduction in leaf water content per leaf dry mass (*Methods: Hydraulic Physiology*).

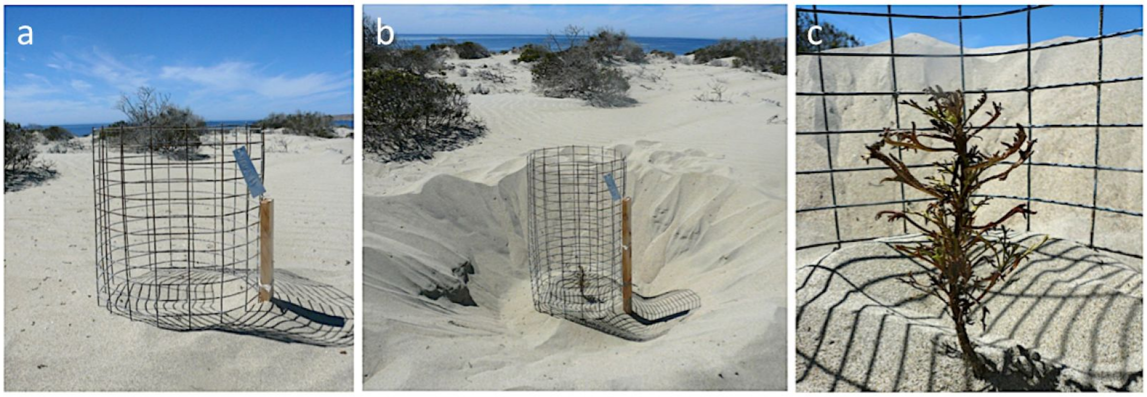

**Figure S7:** Photographs of an experimental hybrid from the desert habitat that was scored as killed due to burial (a) before excavation, (b) after excavation, and (c) close-up after excavation. The plant was not visible above the surface of the sand before excavation, and did not regain turgor pressure after being watered. The total vertical accretion of greater than 20 centimeters of sand occurred over the course of one 48-hour period.

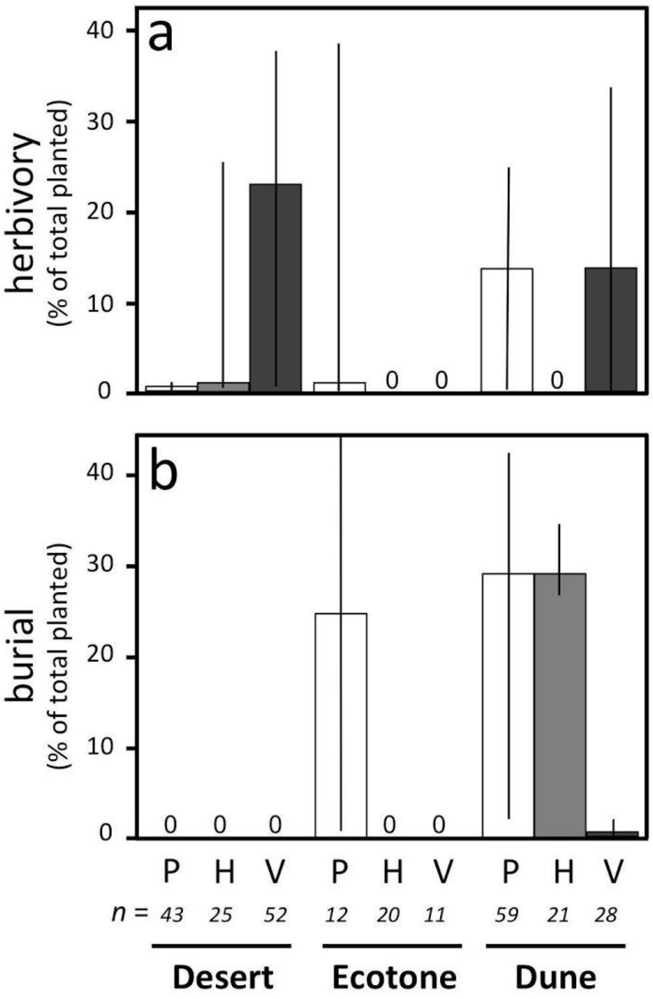

**Figure S8:** Sources of mortality in the reciprocal transplant experiment from (a) herbivory, and (b) burial for *E. palmeri* ("P"), hybrids ("H"), and *E. ventorum* ("V"). Error bars are 95% binomial confidence intervals.

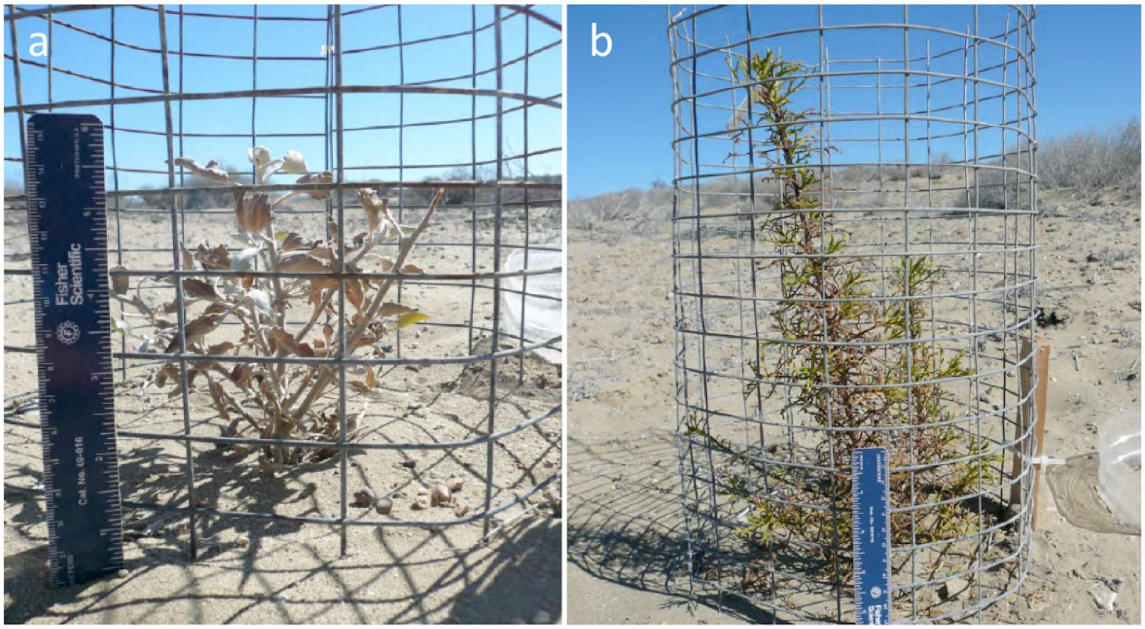

**Figure S9:** Photographs of (a) *Encelia palmeri* and (b) *E. ventorum* from the desert habitat reciprocal transplant experiment, water addition treatment, illustrating the pronounced apical dominance of *E. ventorum* relative to *E. palmeri*. When given supplemental water, *E. ventorum* grew twice as large as the native *E. palmeri* in the desert habitat.

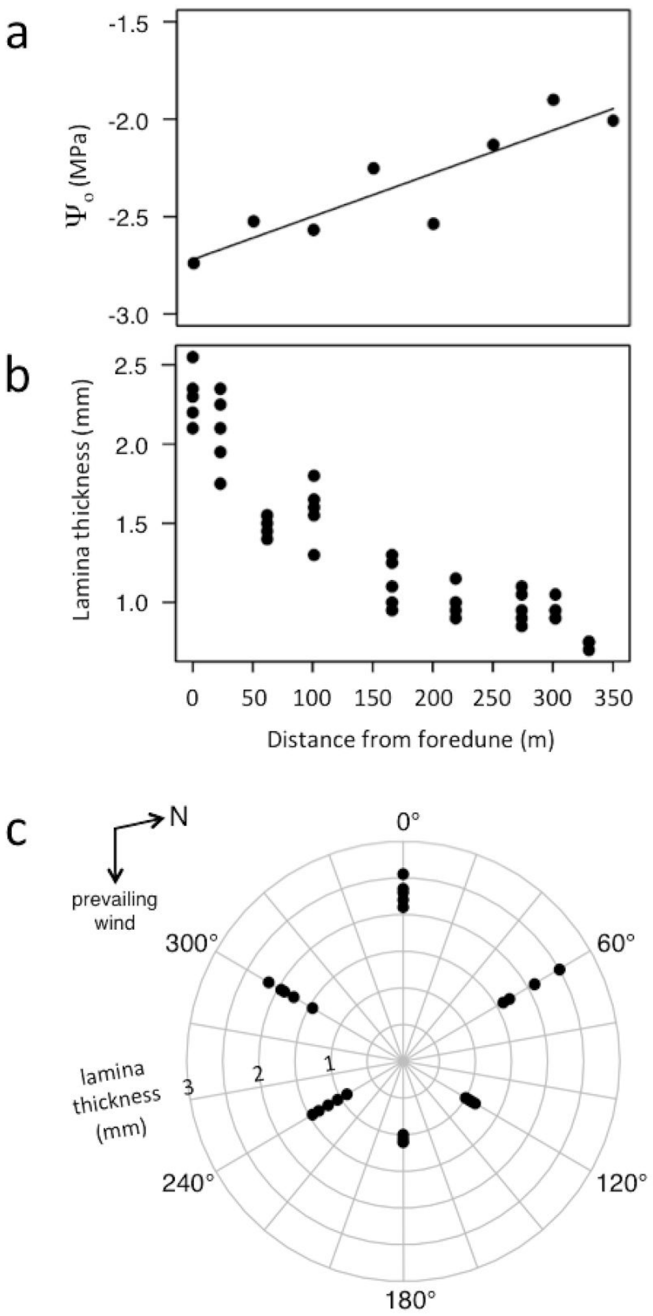

**Figure S10:** (a) Leaf osmotic potentials of *E. ventorum* plants sampled along a transect running from foredune to desert habitats. (b) Leaf thickness used as a proxy for succulence and measured along the same foredune-desert transect. (c) Leaf thickness measured around the circumference of a single exposed foredune plant.

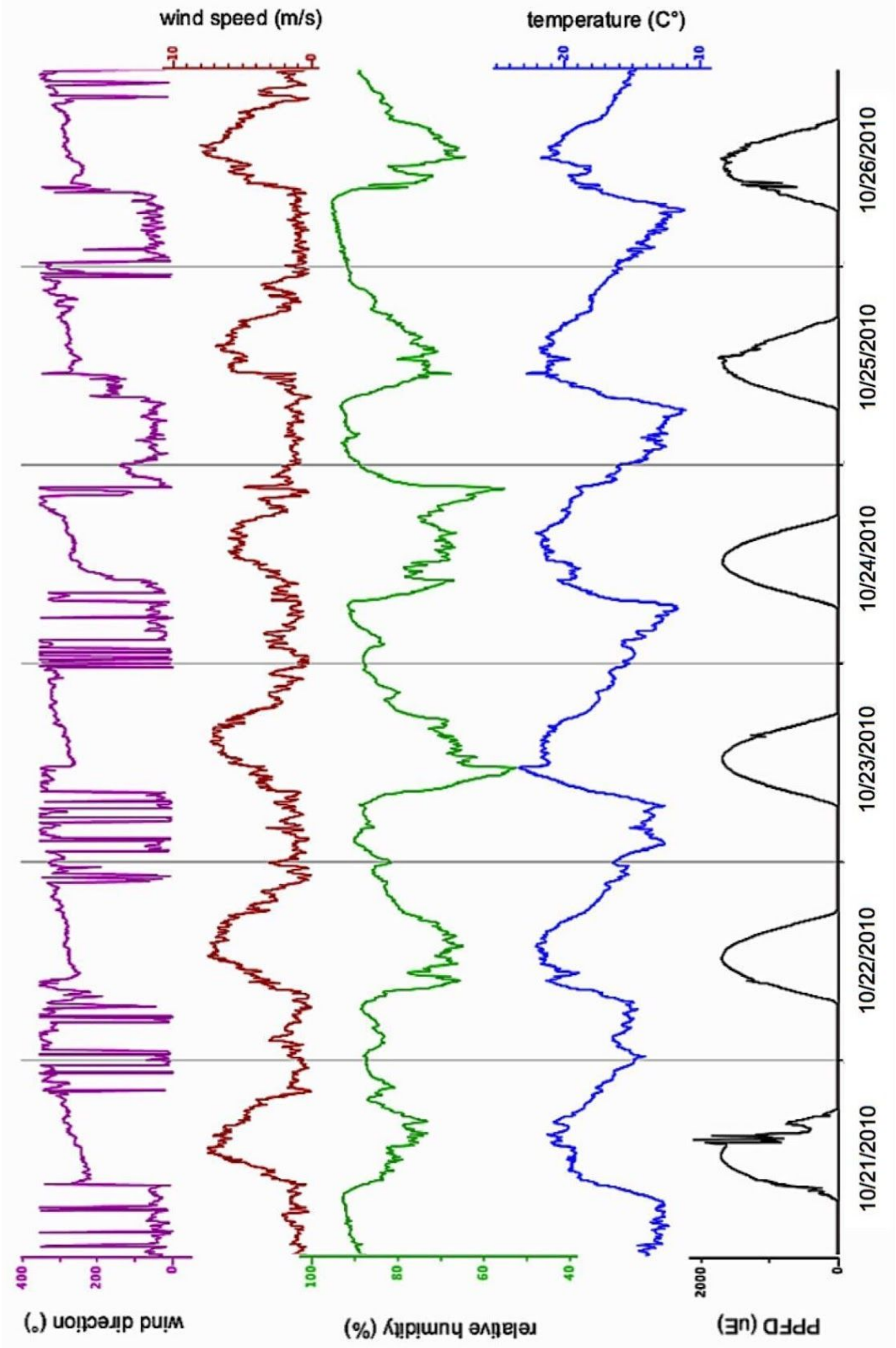

**Figure S11:** Six days of weather from the dune habitat weather station in San Roque during October 2010, the month of planting of the reciprocal transplant experiment. Shown is wind direction (purple), wind speed (red), relative humidity (green), temperature (blue), and photosynthetic photon flux density (black). Afternoon wind direction is consistently onshore from approximately 300°N.

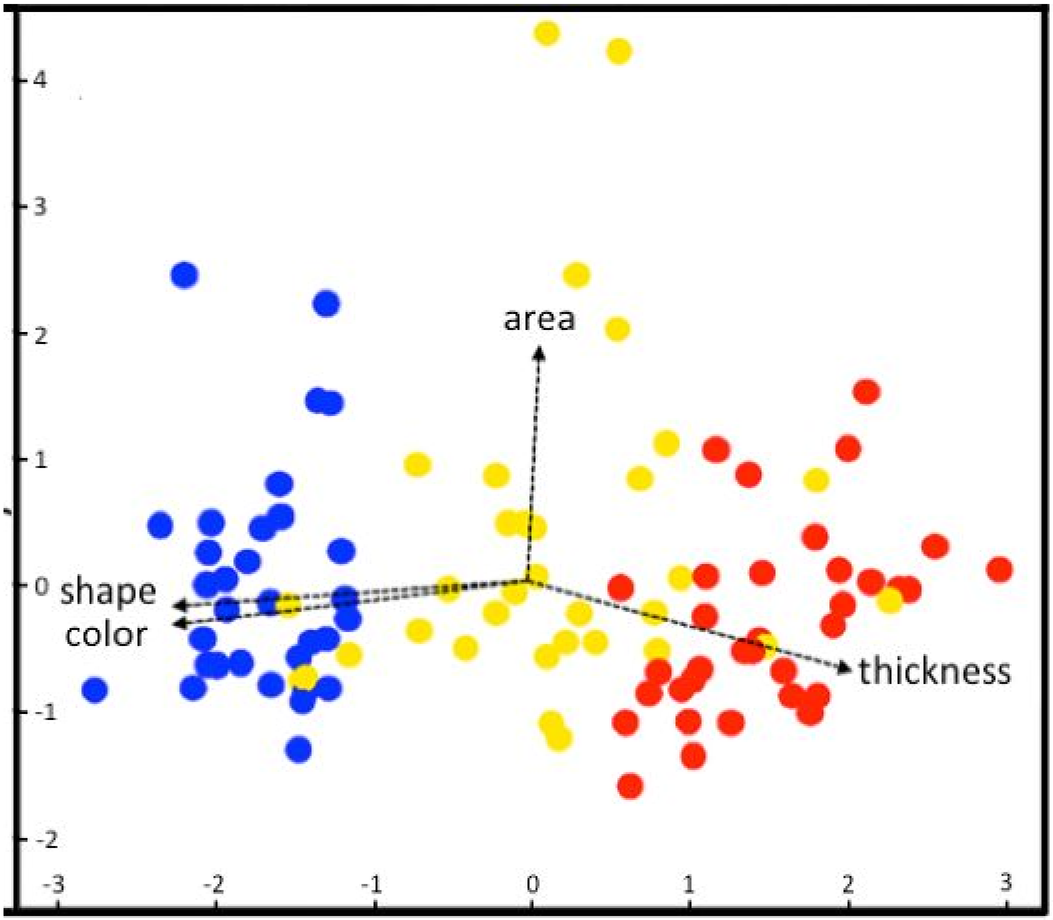

**Figure S12**: First two axes of a principal components analysis of leaf area, shape, color, and thickness measured on individual plants used in the reciprocal transplant experiment: *E. palmeri* (blue), *E. ventorum* (red) and hybrids (yellow). Hybrids have a range of phenotypes, suggesting that the hybrid plants used include later-stage backcrosses.

**Table S1:** Models and AIC scores from analyses of growth, survival, relative fitness, water addition, burial, and herbivory. Models were selected in backwards stepwise fashion, with the saturated model listed first, and the lowest AIC score for that analysis denoted in bold. Not all possible models are shown. GLM indicates general linear model analysis and LL indicates log linear analysis for contingency tables. For LL analysis the dependent variable (survival) references the independent variables with an "×" instead of a "~" to reflect the differences in how the models are specified in R. Statistically significant models at α = 0.05 are denoted with (§), using either analysis of deviance (for GLM analysis) or likelihood ratio tests (for LL analysis), performed on the two models with the highest AIC scores.

| Model | AIC |
|---|---|
| ***Growth*** (GLM) | |
| ln(1+biomass) ~ habitat × phenotype § | **246.40** |
| ln(1+biomass) ~ habitat + phenotype | 254.90 |
| ***Survival*** (LL) | |
| survival × habitat × phenotype § | **36.00** |
| survival × habitat + survival × phenotype + habitat × phenotype | 64.85 |
| survival × habitat + survival × phenotype | 73.71 |
| survival × habitat + phenotype | 82.92 |
| survival × habitat | 89.56 |
| survival × phenotype + habitat | 92.54 |
| survival + phenotype + habitat | 101.75 |
| survival + habitat | 108.39 |
| survival + phenotype | 118.86 |
| ***Relative Fitness*** (GLM) | |
| fitness ~ habitat × phenotype § | **341.60** |
| fitness ~ habitat + phenotype | 354.00 |
| fitness ~ habitat | 358.00 |
| fitness ~ phenotype | 393.1 |
| ***Watering Experiment*** (GLM) | |
| *Desert Habitat* | |
| fitness ~ phenotype × water § | **221.50** |
| fitness ~ phenotype + water | 227.00 |
| fitness ~ phenotype | 254.80 |

| | |
|---|---|
| *Dune habitat* | |
| fitness ~ phenotype × water | 42.59 |
| fitness ~ phenotype + water | 40.64 |
| fitness ~ phenotype | **38.76** |
| ***Burial*** (LL) | |
| burial × phenotype × habitat | 36.00 |
| burial × phenotype × habitat - burial : phenotype: habitat | **34.50** |
| burial × habitat + burial × phenotype | 43.20 |
| burial × phenotype | 73.20 |
| ***Herbivory*** (LL) | |
| *Desert Habitat* | |
| herbivory × phenotype × habitat § | **36.00** |
| herbivory × phenotype × habitat - herbivory : phenotype: habitat | 38.80 |
| *Dune Habitat* | |
| herbivory × phenotype + herbivory × phenotype § | **51.04** |
| herbivory × habitat | 56.83 |

**Table S2:** Hydraulic traits measured, including *a priori* predicted directionality. Error estimates for leaf hydraulic capacitance are 95% confidence intervals, all others ± standard error.

| trait | value | predicted order |
|---|---|---|
| **Mid-day water potential (MPa)** * | | |
| *E. palmeri* | -6.5 ± 2.9 | most negative |
| hybrids | -4.1 ± 1.0 | intermediate |
| *E. ventorum* | -3.9 ± 1.4 | least negative |
| **Leaf hydraulic capacitance (mol kg$^{-1}$ MPa$^{-1}$)** § | | |
| *E. palmeri* | 40.0 (29.8-50.2) | lowest |
| hybrids | 45.9 (36.1-55.6) | intermediate |
| *E. ventorum* | 63.5 (45.4-81.6) | highest |
| **Stem hydraulic conductance (x10$^{-10}$ kg s$^{-1}$ MPa$^{-1}$ m$^{-2}$)** * | | |
| *E. palmeri* | 6.96 ± 0.92 | lower |
| *E. ventorum* | 1.03 ± 0.14 | higher |
| **Xylem vessel size (μm$^2$)** * | | |
| *E. palmeri* | 450.92 ± 11.86 | smallest |
| hybrids | 445.26 ± 13.64 | intermediate |
| *E. ventorum* | 546.46 ± 17.64 | largest |

* denotes significant at α=0.5
§ each value is a slope and so no statistical analysis was performed

**Table S3:** Empirical mass attenuation coefficients (MACs) for unknown crystals in pith tissues and resin ducts, and theoretical MACs for candidate compounds.

| compound | abundance[§] | mass attenuation coefficient ($cm^{-1}$) |
|---|---|---|
| ***Pith crystals*** (empirical) | | |
| *E. palmeri* | 170.80 ± 7.41 | 5.17 ± 0.15 |
| hybrids | 90.25 ± 4.36 | 5.32 ± 0.11 |
| *E. ventorum* | 9.81 ± 3.95 | 5.19 ± 0.11 |
| ***Resin duct crystals*** (empirical) | | |
| *E. palmeri* | 5.5 ± 2.3 | 7.51 ± 0.26 |
| hybrids | 21.2 ± 2.4 | 7.16 ± 0.25 |
| *E. ventorum* | 40.0 ± 5.7 | 7.39 ± 0.20 |
| ***Candidate compounds*** (theoretical) | | |
| $CaC_2O_4$ | na | 5.17 |
| $SiO_2$ | na | 2.73 |
| benzopyrans | na | ~0.2 |
| NaCl | na | 6.26 |
| $MgCl_2$ | na | 7.95 |
| KCl | na | 10.12 |

[§] pith crystal abundance is number per mm of cross sectional stem area, resin duct crystal abundance is number per mm length of stem

**Table S4:** List of all perennial vascular plant species encountered during community sampling and their habitat associations. Data illustrated in Figure 3d.

| species | family | habitat association |
|---|---|---|
| *Jatropha cinerea* | Euphorbiaceae | desert |
| *Fouquieria diguetii* | Fouquieriaceae | desert |
| *Frankenia palmeri* | Frankeniaceae | desert |
| *Lycium andersonii* | Solanaceae | desert |
| *Funastrum arenarium* | Apocynaceae | dune |
| *Helianthus niveus* | Asteraceae | dune |
| *Atriplex julacea* | Chenopodiaceae | dune |
| *Euphorbia misera* | Euphorbiaceae | dune |
| *Stillingia linearifolia* | Euphorbiaceae | dune |
| *Astragalus magdalenae* | Fabaceae | dune |
| *Errazurizia benthamii* | Fabaceae | dune |
